# History shapes regulatory and evolutionary responses to tigecycline in strains of *Acinetobacter baumannii* from the pre- and post-antibiotic eras

**DOI:** 10.1101/2025.01.22.634413

**Authors:** Alecia B Rokes, Alfonso Santos-Lopez, Vaughn S Cooper

## Abstract

Evolutionary history encompasses genetic and phenotypic bacterial differences, but the extent to which history influences drug response and antimicrobial resistance (AMR) adaptation is unclear. Historical contingencies arise when elements from an organism’s past leave lasting effects on the genome, altering the paths available for adaptation. We utilize strains isolated before and after widespread antibiotic use to study the impact of deep historical differences shaped by decades of evolution in varying antibiotic and host pressures. We evaluated these effects by comparing immediate and adaptive responses of two strains of *Acinetobacter baumannii* to the last-resort antibiotic, tigecycline (TGC). When grown in subinhibitory TGC, the two strains demonstrated divergent transcriptional responses suggesting that baseline transcript levels may dictate global responses to drug and their subsequent evolutionary trajectories. Experimental evolution in TGC revealed clear differences in population-genetic dynamics – with hard sweeps in populations founded by one strain and no mutations reaching fixation in the other strain. Transcriptomes of evolved populations no longer showed signatures of drug response, as was seen in the ancestors, suggesting that genetic adaptation may outweigh preexisting differences in transcriptional networks. Genetically, AMR was acquired through predictable mechanisms of increased efflux and drug target modification; however, the two strains adapted by mutations in different efflux regulators. Fitness tradeoffs of AMR were only observed in lineages evolved from the pre-antibiotic era strain, suggesting that decades of adaptation to antibiotics resulted in preexisting compensatory mechanisms in the more contemporary isolate, an important example of a beneficial effect of historical contingencies.

**SIGNIFICANCE STATEMENT:** *Acinetobacter baumannii* is a high priority pathogen often causing multidrug resistant nosocomial infections. Many healthcare systems experience clonal outbreaks of *A. baumannii* infections, yet treatment strategies are often strain-agnostic, ignoring the importance of strain differences. We show that historical differences between two strains, one isolated prior to widespread antibiotic use and the other following decades of selection to clinical conditions, dictate transcriptional patterns and response to a last-resort antibiotic and influence the genetic and phenotypic routes of resistance adaptation. While our study focuses on two reference strains of *A. baumannii,* these findings can be more broadly applicable to other pathogenic organisms in which a better understanding of the forces influencing resistance adaptation is essential for combating the antimicrobial resistance crisis.

## INTRODUCTION

Evolution is an inescapable process, as the worsening antimicrobial resistance (AMR) crisis demonstrates (1). However, the AMR crisis will be less daunting if we are better able to understand, predict, and direct the evolutionary outcomes of antibiotic treatment (2, 3). With better resolution of factors affecting the routes to resistance, we may be able to work toward directing evolution to avenues with exploitable resistance tradeoffs (4, 5). However, genotypic and phenotypic routes to resistance, as well as the resulting fitness tradeoffs, may be highly influenced by strain history, decreasing the range and applicability of evolutionary strategies to counteract the evolution of AMR (6, 7).

There are three primary forces, often intertwined, that influence evolution: history, chance, and selection (5, 8, 9). Natural selection causes the most fit alleles in a population to rise in frequency while less fit mutants are lost from the population. Selection needs heritable phenotypic variance to act upon that is produced by mutations arising randomly, a factor of chance. Chance events also occur in the demography of microbial populations, such as population bottlenecks arising during infections (10). The outcome of a mutation is also affected by the genetic background in which it arises (11, 12), combining chance and the final force of evolutionary history, which can influence adaptation in diverse ways but remains understudied.

Evolutionary history encompasses the genetic and phenotypic remnants from a bacterium’s past including biotic and abiotic factors, such as interactions with other cells and previous environmental exposures like antibiotics, respectively (13). The phylogenetic origin of the bacterium is also a major component of evolutionary history. These differences can impose historical contingencies through which preexisting elements in the genome, transcriptome, or phenome alter or constrain evolutionary paths available for adaptation (13, 14). History may dictate the type, identity, rate, and order of mutations acquired (15–18). Additionally, history likely influences fitness and phenotype associated with a mutation, thereby dictating evolutionary trajectories and outcomes (19–21). This study seeks to understand how historical contingencies influence evolutionary trajectories under the strong selective pressure of antibiotic treatment (22).

Contingencies imposed by history exacerbate our inability to accurately predict bacterial responses to antibiotics and AMR evolution. *Acinetobacter baumannii* is a nosocomial pathogen of increasing concern due to limited and often ineffective treatment options (23). With time and increased antibiotic use, strains of *A. baumannii* have acquired mutations and mobile genetic elements, on top of many conserved genes, that endow desiccation resistance, stress tolerance, and drug resistance (24, 25). Such historical differences could cause archaic strains, which are often represented by laboratory reference strains, to respond and evolve differently than contemporary infectious strains (26, 27). Importantly, hospitals typically experience clonal outbreaks of *A. baumannii* caused by closely related isolates of the same sequence type (28, 29). Across hospital systems, *A. baumannii* infections exhibit high diversity (30, 31), but only a handful of treatments are used. This practice ignores the importance of genetic background and evolutionary history in the treatment process, likely leading to differences in treatment response (32, 33). A better understanding of how various strains respond and adapt to antibiotics would result in improved patient outcomes.

The widespread use of antibiotics in clinical practice has dramatically affected pathogen evolution (7, 34). Comparing strains isolated before and after the clinical introduction of antibiotics provides an excellent framework to study how deep historical differences influences the evolution of resistance (35). To this end, we used two strains of *A. baumannii,* one that was isolated early in the antibiotic era and has not been exposed to many contemporary antibiotics, and one that is better representative of current multi-drug-resistant (MDR) clinical infections having been isolated much more recently. Comparing these strains enables assessment of how a history of evolving with the human host and novel antimicrobial pressures impacts resistance evolvability.

Resistance adaptation can be investigated at multiple levels: from the narrowest, the site-specificity of individual mutations, to more systematic resistance pathways, and even more broadly, at the level of population-wide responses and adaptive dynamics (36, 37). This study looks within and across these levels to determine the areas in which evolutionary history is most influential. We find that deep history influences the basal transcriptome, driving differences in the response to TGC. The strains also differed in their evolved responses to prolonged TGC stress at both narrow molecular-genetic and broader population-genetic levels, despite superficial similarities. This work highlights the importance of evolutionary history in the response and adaptation of a high-priority pathogen to a last-resort antibiotic, showing that strain differences can reduce the predictability of treatment outcomes.

## RESULTS

### Genomic and phenotypic classifications of the strains

Previous studies of historical differences in strain response to antibiotics have used strains separated by relatively few mutations (9). Here, we study effects of deeper history on the evolution of AMR in two different *A. baumannii* strains isolated nearly 60 years apart, *i.e.* before and after the widespread clinical use of antibiotics. The first strain, 17978UN, is a variant of the commonly used laboratory reference strain ATCC 17978 (38). This strain was isolated in 1951 in the early stages of the introduction of antibiotics in clinical practice (39). In contrast, the second strain, AB5075-UW, is a variant of AB5075, which was isolated in 2008 from a MDR infection representative of the current global clinical burden of *A. baumannii* (40, 41). Therefore, the evolutionary histories of both strains have likely been shaped by different exposures to antibiotics, hospitals, and the human host. In many ways 17978UN represents archaic *A. baumannii* infections in comparison to contemporary MDR infections represented by AB5075-UW.

We first characterized genomic and phenotypic differences between the two strains to evaluate their historical distinctions. 17978UN is a member of sequence type ST437 whereas AB5075-UW is assigned to ST1, part of the clinically dominant clonal complex 1 (40). Their genomes share an average nucleotide identity of 97.5% and 3265 homologous core genes. 17978UN contains an additional 694 accessory genes whereas AB5075-UW encodes 751 accessory genes (Data S1), with no significant differences in gene content grouped by Clusters of Orthologous Genes (COG) categories (Table S1). However, as expected by its lineage and history of antibiotic exposure, AB5075-UW has significantly more resistance-associated elements than 17978UN (Table S1) and is more resistant to a variety of antibiotics (Table S2). Notably, both strains were susceptible to at least two different classes of antibiotics.

These significant differences in genome content and existing AMR led to the hypothesis that the strains would respond differently to a contemporary, clinically relevant antibiotic. Tigecycline (TGC) is a translation-inhibiting antimicrobial compound that was approved for clinical use in 2005 (42). Resistance to TGC in clinical isolates of *A. baumannii* is typically attributed to overexpression of efflux pumps or ribosomal modifications (43, 44). The minimum inhibitory concentration (MIC) of TGC differed only slightly between the strains (17978UN MIC_TGC_ = 0.125 µg/mL, AB5075-UW MIC_TGC_ = 0.25 µg/mL). Further, subinhibitory TGC (0.06 µg/mL) imposed approximately 75% and 60% fitness defects in 17978UN and AB5075-UW, respectively (Fig. S1). This comparable level of growth inhibition by TGC enables the study of influences of deep evolutionary history on both transcriptional response and subsequent evolutionary adaptation.

### Strain-dependent transcriptional response acts on historically differentially utilized genes

Given the stress imposed on growth by TGC, even at subinhibitory levels, we predicted that transcriptional responses to this drug would be largely conserved between strains, likely consisting of stress-response genes, drug resistance mechanisms, or ribosomal genes specific to the mechanism of action of TGC (45, 46). We compared transcript levels following growth in subinhibitory TGC to growth in minimal media lacking antibiotic. The defined minimal media used here and throughout was M9 salts buffered supplemented with glucose, amino acids, and other elements as described in the methods (47). As expected, TGC stress caused large transcriptional changes biased toward gene downregulation in both strains, but there was surprisingly minimal overlap among differentially expressed genes between strains (Fig. 1 *A* and *B*). Only 144 genes, representing 24% of the total downregulated genes, and 53 genes, representing 18% of upregulated genes, were differentially expressed in the same direction in both strains. One might intuit that the limited shared response to drug may have resulted from transcriptional differences among strain-specific genes, but this was not the case. Rather, most genes that were differentially expressed were within the shared genome (Fig. 1 *A* and *B*; Data S1), including many genes with the greatest changes in expression that shifted in opposite directions between strains (Fig. S2 and Fig. S3). Together these results show that the response to TGC stress is historically contingent.

**Fig. 1:**
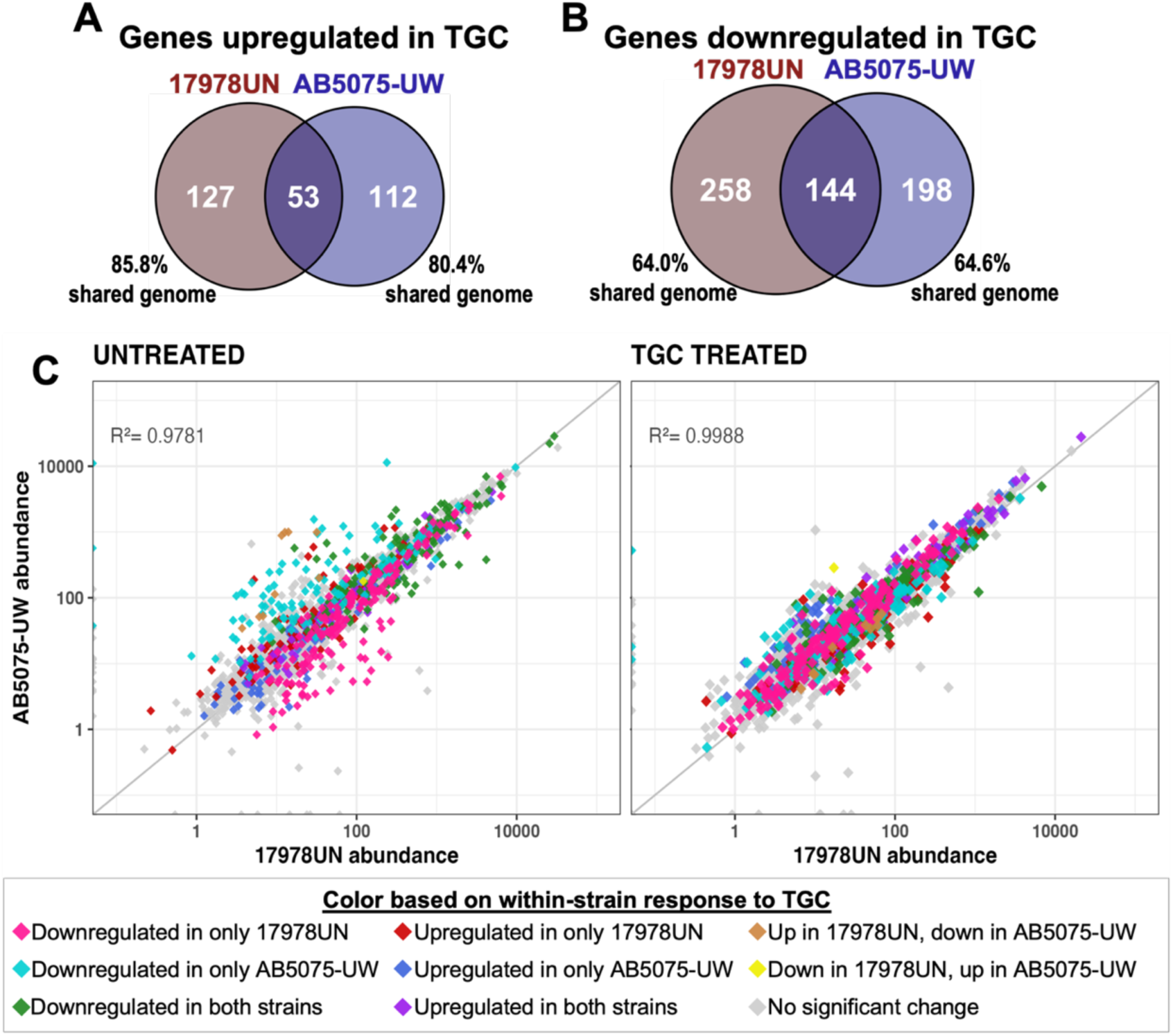
Transcriptional response to TGC is highly strain dependent and results in a shift toward a more conserved transcriptional state. Counts of differentially expressed genes, upregulated (*A*) or downregulated (*B*), in each strain in response to TGC treatment. Overlapping regions represent genes that were similarly differentially expressed in both strains. Percentage of genes that were significantly differentially expressed in one strain but are still encoded by both strains (shared genome) is indicated, showing minimal involvement of accessory genes. (*C*) Each point represents the abundance of one gene in each strain in that condition (left: untreated plain M9plus media; right: TGC treated with 0.06 µg/mL TGC). Diagonal y = x line represents equal gene abundance in both strains and R^2^ linear regression fits to this line are indicated. Points are colored based on within-strain significant differential expression upon TGC treatment. See Animation S1 for an animation of these transcriptional responses, and Fig. S4 for accessory gene transcriptomic responses and a version of this plot with colors separated.

To test whether baseline transcriptional differences explained varied strain response to TGC, we compared transcript abundance of conserved genes in the absence of drug (Fig. 1*C*, *left*). This comparison revealed differences in baseline transcriptional state that, upon TGC pressure, caused expression to converge upon a common pattern of gene usage (Fig. 1*C*, *right*). Genes with higher basal transcription in each strain were preferentially downregulated within that strain in response to TGC treatment (Fig. 1*C*; Fig. S4), producing a more conserved transcriptional state under TGC pressure (Animation S1). These historical differences in basal gene expression explain the strain-dependent responses to TGC pressure.

### Experimental evolution for tigecycline resistance results in strain-dependent tradeoffs between fitness and resistance

Given the genomic signatures of history seen in the transcriptional response to TGC, we hypothesized that evolutionary adaptation to this antibiotic would proceed along different paths in each strain. We propagated three replicate populations of both strains for 12 days in increasing concentrations of TGC (methods and Fig. S5). All populations evolved excess resistance beyond the drug concentration in the media, which was maintained throughout the experiment, such that all populations became nearly five times more resistant to TGC than their respective ancestors (Fig. 2).

**Fig. 2:**
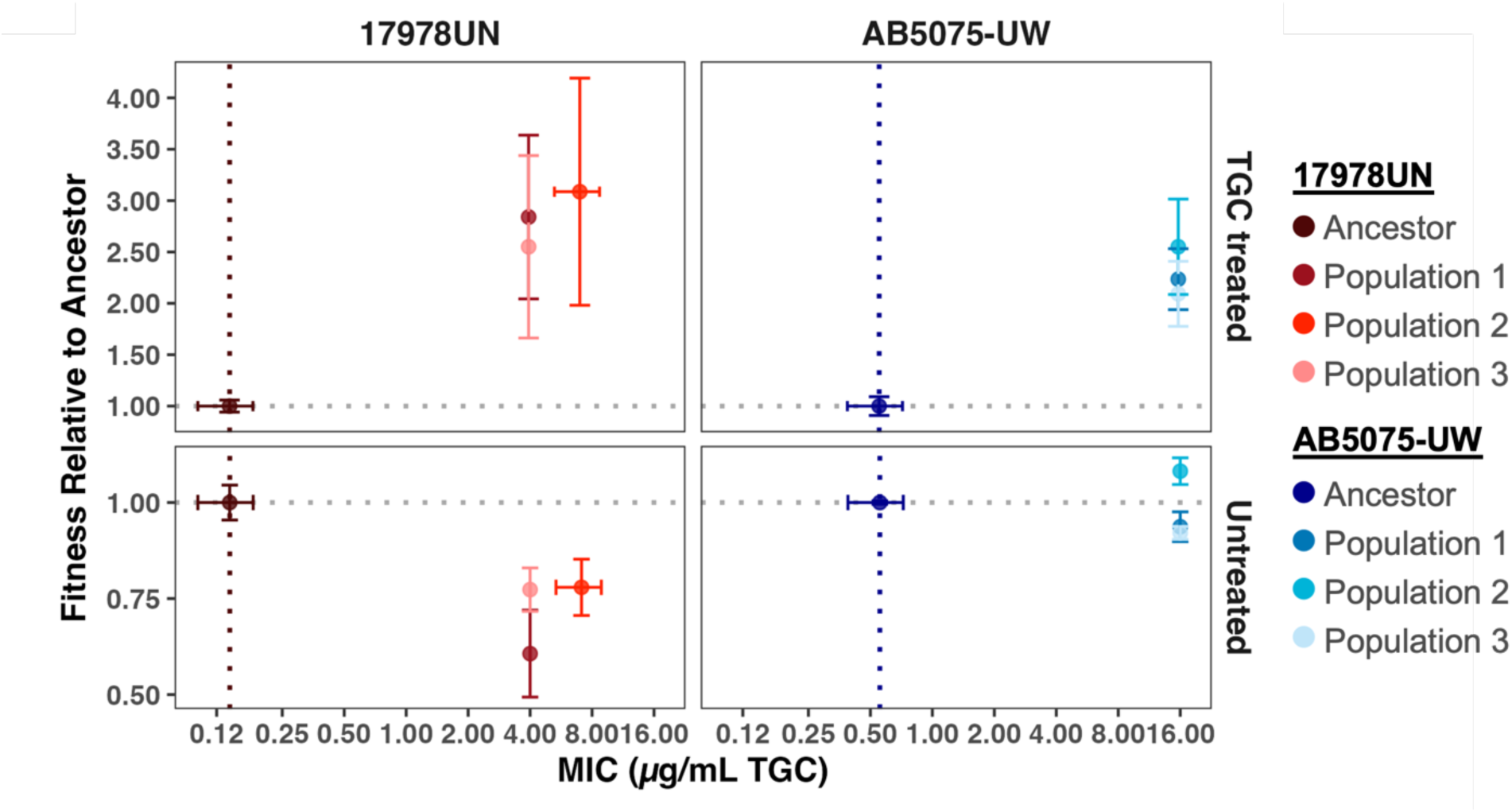
Strain specific patterns in fitness-resistance tradeoffs. Minimum inhibitory concentration (MIC) to TGC was measured for ancestors and for day-12 TGC-evolved populations. Relative fitness was calculated from growth curves as described in the methods. Fitness of 1.00 (gray horizontal dotted line) indicates the population grew equally well to its respective ancestor in that environment (untreated M9plus or TGC treatment at 0.06 µg/mL TGC). Points represent mean across replicates and error bars show standard deviation of the mean.

Antibiotic resistance is often associated with a fitness cost and these costs can depend on the strain background (17, 48–50). We expected to find fitness deficits of TGC-evolved populations when grown in the absence of drug, whereas we expected higher fitness of TGC-evolved populations than their respective ancestors when grown in subinhibitory TGC. As expected, both sets of evolved populations exhibited substantial fitness increases in subinhibitory TGC. However, evolved fitness responses in the absence of drug differed. Populations derived from the laboratory reference strain, 17978UN, incurred a 25-50% reduction in fitness, whereas no measurable fitness cost was observed for populations derived from the more recently isolated clinical strain, AB5075-UW (Fig. 2). These results show that evolutionary tradeoffs between fitness and resistance to TGC differ by founding strain; and suggest that contemporary MDR strains may adapt to new drugs with limited collateral fitness costs.

### Varying population dynamics within conserved mechanisms of TGC resistance

The divergent relationships between fitness and resistance indicated that the genetic pathways to TGC resistance may differ between the two strains. We used longitudinal whole-population, whole-genome sequencing to investigate the population-genetic dynamics of adaptation. Following prior evolution experiments in antibiotics (47, 51), we predicted that antibiotic pressure in well-mixed populations would be sufficiently strong that mutations providing resistance should rapidly sweep within the population. Indeed, mutations in genes associated with drug resistance rapidly fixed in all populations of AB5075-UW (Fig. 3; Data S2), but surprisingly, no mutations reached fixation in populations of 17978UN. A closer inspection of the mutations in each 17978UN line suggests that in total, they represent a soft sweep of resistance-associated mutations. Soft sweeps occur when multiple independent sub-lineages acquire mutations in the same genes; individual mutations never reach fixation, but the sum of all co-existing analogous mutations in a gene or pathway approach fixation within the population (52).

**Fig. 3:**
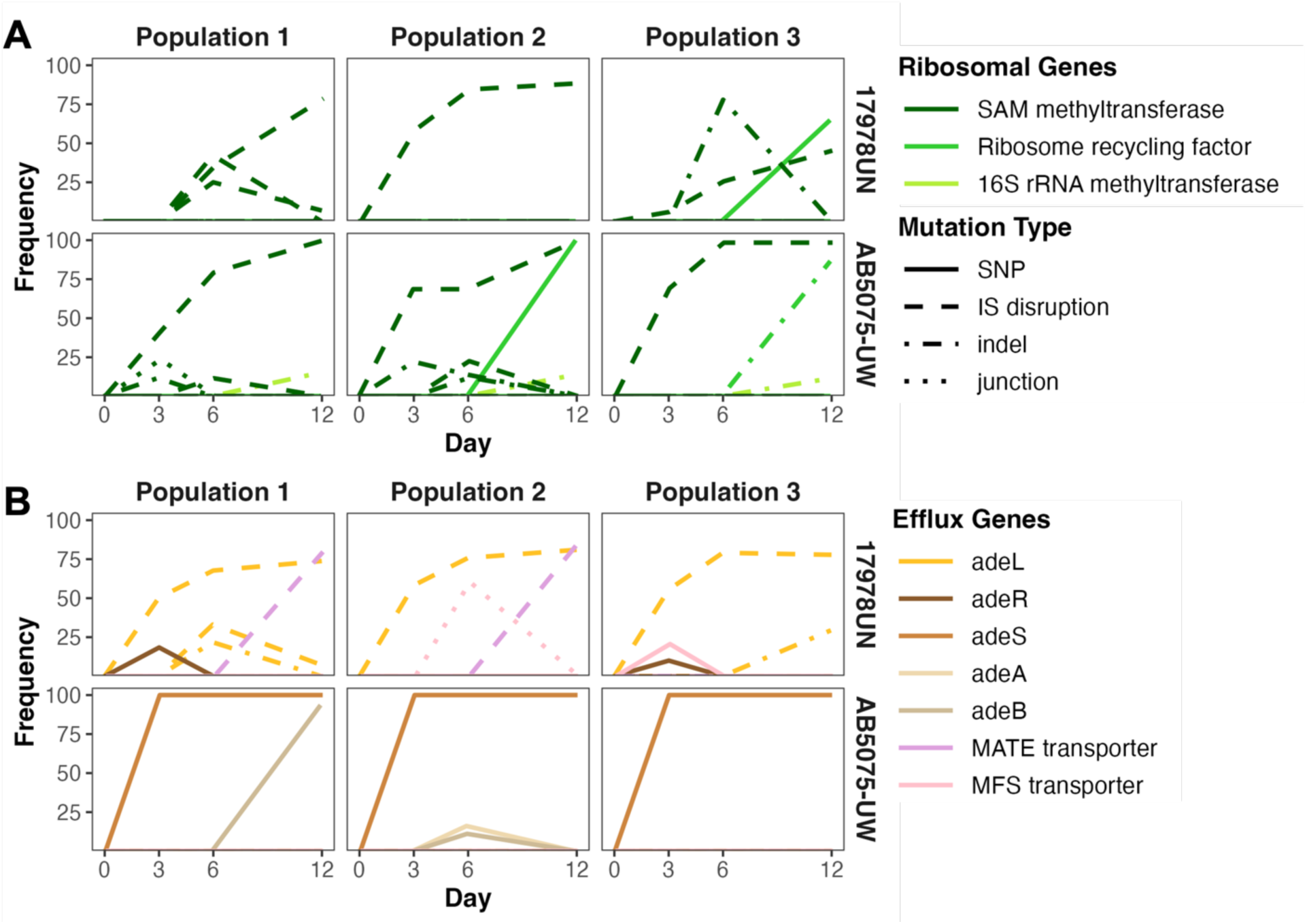
Whole-population genome sequencing reveals similar resistance mechanisms but different population dynamics between strains. Evolved mutation frequencies over time, with line colors corresponding to affected gene and line patterns indicating mutation type. Abbreviations: SNP = single nucleotide polymorphism; IS = insertion sequence, indel = insertion or deletion, and junction = structural variant indicated by a new junction call. Multiple lines of the same color indicate more than one unique mutation arose in that gene. (*A*) Mutations in genes associated with the ribosome. (*B*) Mutations in genes associated with efflux pumps.

The evolved mutations pointed to two primary mechanisms of resistance to TGC: modification of the enzymatic target of the drug, the ribosome, and decreasing the intracellular concentration of antibiotic, primarily through efflux pumps. All replicate populations of both strains featured high-frequency mutations affecting the same pathways for drug resistance and even in the same gene. This gene encodes a *S*-Adenosyl-methionine-dependent methyltransferase (SAM-MT; Fig. 3*A*) that has been associated with creating methylation patterns required for TGC binding to the ribosome (53). Interestingly, all SAM-MT mutations that reached high frequency under TGC stress were gene disruptions caused by mobilized insertion sequences that are predicted to eliminate function (Data S2). Although each population acquired mutations in the same SAM-MT, the specific insertion sequence causing the loss of function as well as the site of insertion varied among lineages, with more variability within populations of AB5075-UW. In addition to the SAM-MT, other genes related to the target of TGC acquired mutations, such as ribosome recycling factors (Fig. 3*A*; Data S2) that likely aid in rescuing stalled ribosomes (43).

All populations also evolved mutations in regulators of efflux pumps, but the specific resistance-nodulation-division (RND) pump that was affected differed between strains. Populations of 17978UN also acquired mutations affecting other systems predicted to reduce intracellular concentrations of TGC. For instance, two 17978UN populations acquired mutations in major facilitator superfamily (MFS) transporters that rise to high frequency (Fig. 3*B*; Data S2) (54). In summary, the identities, timing, and frequencies of evolved mutations differed between strains, whereas replicate populations founded by the same strain exhibited high gene-level parallelism.

### Strain-dependent efflux pump preferences

All populations adapted to TGC through mutations in drug efflux systems (Fig. 3*B*), yet different strains evolved with different genetic signatures. *A. baumannii* encodes three RND-efflux pumps known to be associated with drug resistance adaptation (55, 56). Two-component system *adeRS* regulates efflux operon *adeABC,* while the other two pumps, *adeFGH* and *adeIJK* are regulated by negative regulators *adeL* and *adeN,* respectively (Fig. 4*A*) (57). Both strains encode all three pumps with a high level (97-100%) of amino acid conservation (Table S2), with the only exception that 17978UN does not encode the outer membrane protein, AdeC, but AdeK can complement function (55). Populations of AB5075-UW acquired nonsynonymous mutations throughout the *adeRSABC* efflux system, altering *adeA*, *adeB*, and *adeS*. One of two predominant single nucleotide polymorphisms (SNPs) in *adeS* (R152K and G160S) fixed in all AB5075-UW populations, and an additional SNP in *adeB* (E939Q) was nearly fixed in one population (Data S2). In contrast, each evolved 17978UN population acquired mutations in *adeL* caused by an insertion sequence (IS) disruption in this gene that rose to high frequency (Fig. 3*B*; Data S2). This disruption exhibited site-specific parallelism in which the mutation was identical within all three TGC-evolved populations, but never reached 100% frequency. As an example of the soft sweep dynamics described previously, in evolved population 3 we observe an additional mutation in *adeL* and the two mutations in this gene, cumulatively, reach fixation. Interestingly, a nonsynonymous SNP in *adeR* also arose in 17978UN populations 1 and 3 on day three but it was quickly outcompeted by *adeL* mutations (Fig. 3*B*), further emphasizing the strain-dependence of efflux mutations.

**Fig. 4:**
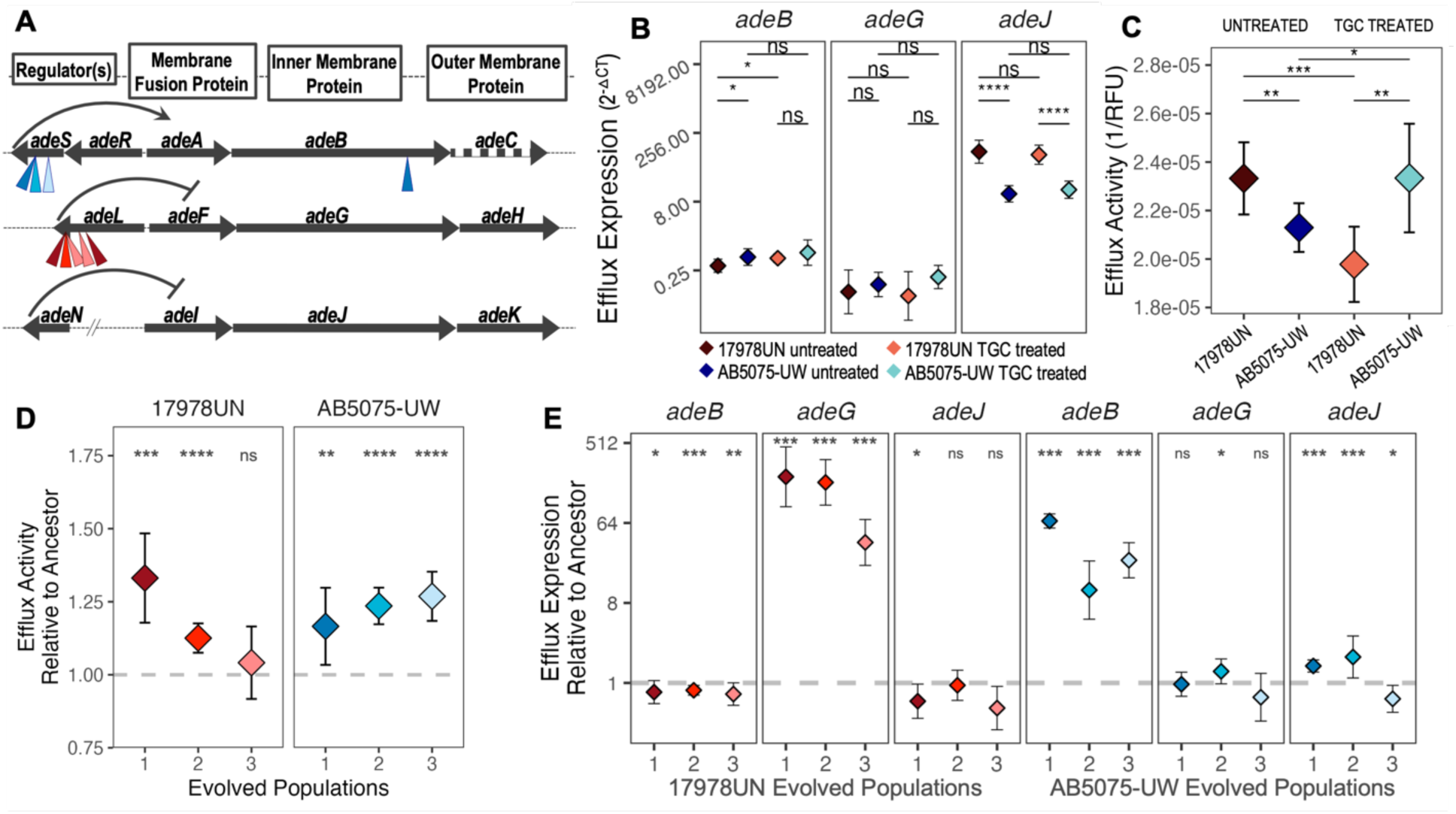
Genetic distinctions of evolved efflux expression and activity. (*A*) Representation of the RND efflux pump operons/regulators. All RND genes are highly conserved between the two strains (supplemental table 2), except that 17978UN does not encode *adeC.* Arrow size is scaled to the length of the gene. Inverted triangles depict mutations present on day 12, colored by the population in which they are found. Ancestral efflux activity (*B*) and expression (*C*) in treated (0.06 µg/mL TGC) or untreated states. (*D*) Efflux activity of the evolved populations when grown in the untreated state relative to respective ancestor. (*E*) Efflux expression in evolved populations normalized by the respective untreated ancestor. Points represent mean and error bars depict standard deviation. Statistically significant differences were determined with t-tests in B and C with the comparisons depicted by lines, and with Wilcoxon tests compared to an ancestral value set at 1 in D and E. Significance criteria: ns for p > 0.05, * for p ≤ 0.05, ** for p ≤ 0.01, *** for p ≤ 0.001, and **** for p ≤ 0.0001.

There are many ways in which historical contingencies could cause such strain-specific preferences of efflux targets. One possibility is that the sequences of these target genes had already differentiated, but all sites that acquired adaptive mutations were initially identical in the ancestral genomes (Table S3). Another possibility is that the ancestral strains had previously evolved differential regulation of these pumps by other mechanisms. To test this notion, we measured expression of efflux genes in the presence and absence of subinhibitory TGC but found comparable levels of expression of *adeABC* and *adeFGH* (Fig. 4*B*), discounting this explanation. We found that 17978UN exhibits relatively greater expression of the *adeIJK* pump in both growth conditions, but this pump was not the target of selection in either strain. In the absence of TGC, 17978UN had increased efflux activity compared to AB5075-UW (Fig. 4*C*), possibly due to the differences in AdeIJK expression. Interestingly, drug pressure induced significantly different states of efflux activity, increasing efflux activity in AB5075-UW but decreasing it in 17978UN (Fig. 4*C*). As expected, there were significant differences in efflux activity and expression in the evolved populations, confirming that the mutations in efflux regulators produced a meaningful phenotype. All evolved populations, except 17978UN evolved population 3 which still trended higher, evolved significantly increased efflux activity than their respective ancestors (Fig. 4*D*). Lastly, we show that the mutated regulators specifically increase expression of their associated pump (Fig. 4*E*). In summary, neither sequence divergence nor functional differences in the primary targets of selection explained the alternative evolutionary pathways to increased drug efflux. These results strongly suggest that more complex interactions between efflux systems and the genome favor different routes to drug adaptation.

### Diminished transcriptional response and historical influences following TGC adaptation

The ancestral strains exhibited divergent transcriptomic changes in response to TGC stress (Fig. 1), but both adapted to TGC via broadly similar mechanisms. To evaluate the transcriptional signature of evolved drug tolerance, we conducted RNA sequencing of evolved populations grown with or without TGC treatment and found remarkably few differences (Fig. 5*A*). Compared to the ancestral response, where many genes changed in abundance upon TGC treatment, all evolved populations exhibited nearly identical transcriptomes in the two conditions. This trend is more conspicuous in populations of AB5075-UW, where no genes are significantly different (p-value < 0.05 and absolute value log fold change > 1) between the two conditions, while a few genes remain significantly differently expressed upon TGC stress in populations of 17978UN (Fig. S6, Data S3).

**Fig. 5:**
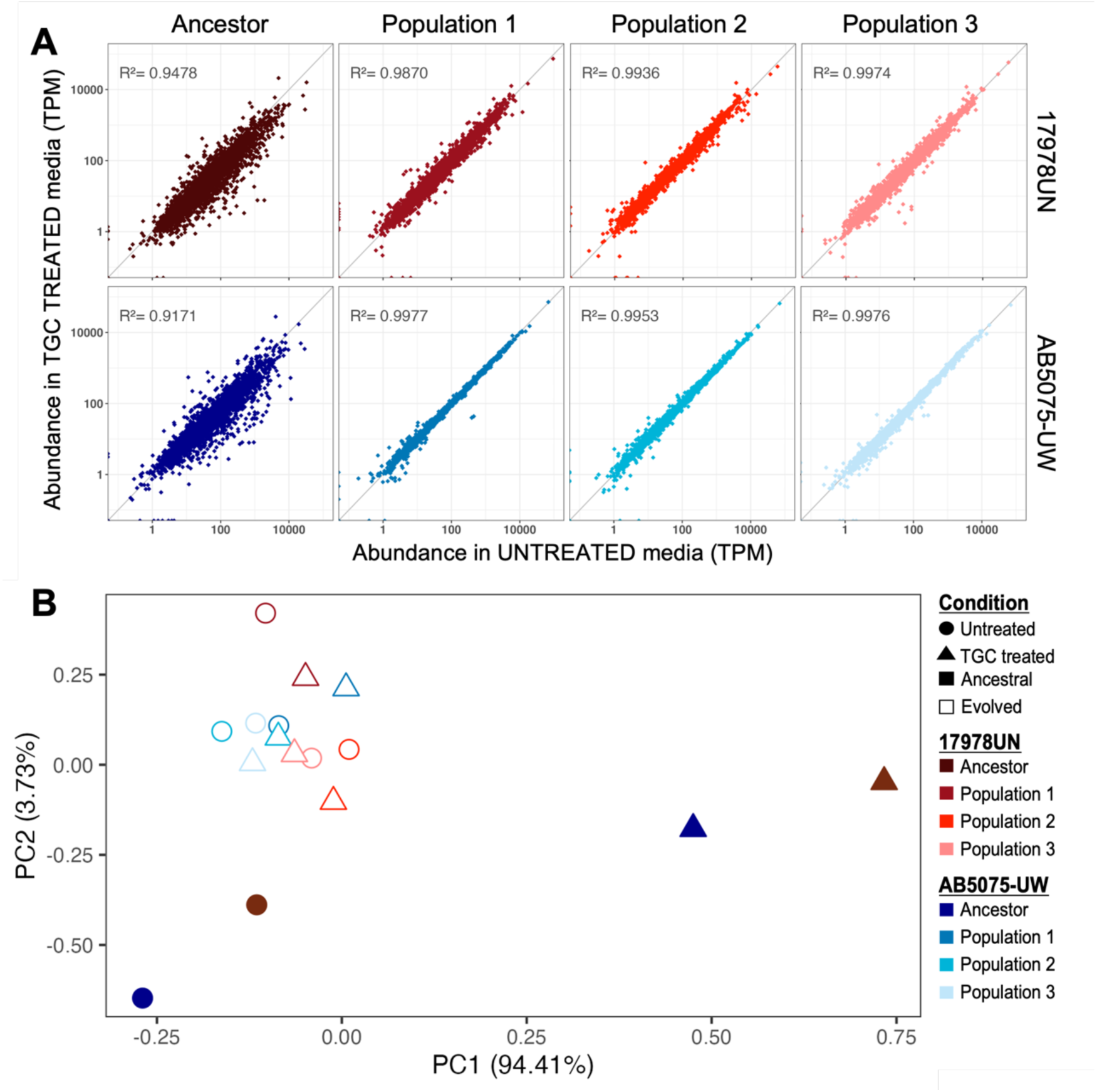
Transcriptional response evolves to minimally react to TGC pressure. (*A*) Conserved transcript abundance regardless of TGC treatment in all evolved populations. Each point represents the abundance of one gene in each sample. Point position depicts expression in each condition (untreated M9plus, or 0.06 µg/mL TGC). Top row shows ancestor and TGC-evolved populations for 17978UN, and the bottom row is the same for AB5075-UW. Diagonal y = x line represents equal gene abundance in both conditions and R^2^ linear regression fits to this line are indicated. (*B*) Principal component analysis showing that ancestral transcriptomes are strain dependent, but TGC treatment produces a conserved shift. Evolved population transcriptional adaptation is conserved between strains and indifferent to TGC treatment. Point color indicates sample, shape indicates culture condition either untreated or 0.06 µg/mL TGC treated, fill indicates either ancestral or evolved populations.

While the muted transcriptional response to drug is observed in all evolved populations, it does not tell us if evolved transcriptional responses are conserved between strains. Principal component analysis of transcript abundance confirmed that TGC treatment shifts the initially diverging ancestral transcriptomes in the same direction while still preserving strain differences (Fig. 5*B*). Transcriptomes of evolved populations, either treated or untreated, strongly grouped together. This cluster of evolved populations groups more closely with the untreated ancestors, further supporting limited transcriptional response to TGC stress by evolved populations (Fig. 5*B*; Fig. S6). Taken together, these convergent evolved transcriptomes show that genomic drug adaptations alleviate the need to transcriptionally respond to TGC pressure and result in reduced historical signatures on the transcriptome compared to the divergent ancestral transcriptomes.

## DISCUSSION

Antimicrobial resistance (AMR) is a problem of evolution that is most often observed in retrospect, when treatment fails. We still have much to learn about how AMR evolves, including the diversity of genetic pathways to resistance (22), the extent to which these pathways depend on the environment (47) including the host immune status (58), and whether these pathways differ among historically divergent strains (18, 20). In this study, we focused on this last factor by evaluating how reference strains isolated nearly 60 years apart, spanning eras of antibiotic usage, differ in their adaptation to a clinically significant drug. Although both strains of *A. baumannii* adapted to grow in media containing tigecycline (TGC) via common pathways, the specific genes and mutations differed with the founding strain. On the other hand, replicate populations derived from the same strain evolved in parallel, indicating a level of genetic predictability within but not between strains.

Perhaps the most striking difference between strains was the fitness consequence of TGC adaptation: lineages derived from the modern clinical strain experienced no measurable fitness costs whereas those derived from the historical strain, 17978UN, incurred deficits of 25% or more (Fig. 2). Fitness effects are often related to the stability or persistence of resistance. Costly resistance has long been proposed as a beneficial trait for humanity that can preserve drug efficacy in the face of selection (3, 59, 60). Here we find an alarming lack of a fitness cost of evolved resistance in lineages derived from the contemporary clinical isolate, AB5075-UW. The evolutionary history of AB5075-UW includes frequent antibiotic exposure prior to our studies that could have selected for compensatory adaptations that alleviate costs of TGC resistance (61). The lack of a fitness-resistance tradeoff in AB5075-UW suggests that passive loss of resistance once drug pressure is alleviated will be rare in related strains, increasing the need for alternative treatment options.

Differences in the genetic causes of resistance between strains offer evidence that historical contingency influences routes to AMR. Many bacterial genomes including *A. baumannii* encode multiple RND efflux pumps and to explain this apparent redundancy, a hierarchical model of sequential activation has been proposed (54, 57, 62). We find that the first, dominant mutations affecting drug efflux in each strain affected regulators of different pumps, with SNPs in *adeS* sweeping to fixation in AB5075-UW populations and gene disruptions caused by IS in *adeL* reaching high frequencies in 17978UN populations (Fig. 3). This suggests that the hierarchy of these pumps differs between strains in the face of TGC pressure and raises the possibility that variation in their regulatory network or gene content evolved in the past. Perhaps the diminished priority of the AdeRSABC pump in 17978UN stems to the lack of *adeC* in this strain (Fig. 4*A*; Data S1). These lineages would need to accumulate two mutations to overexpress one functional pump: one that upregulates AdeAB and one that upregulates AdeK to serve as the outer membrane protein (55). The historical contingency imposed from the loss of *adeC* sometime in the history of 17978UN may be driving the strain-dependent efflux preferences for drug adaptation.

Both the spectra of evolved mutations and their dynamics differed between strains, suggesting intrinsic biases that influence genetic diversity. Despite having the same number of encoded insertion sequences (IS) as AB5075-UW (Table S1), nearly all high-frequency mutations in populations of 17978UN were caused by IS interrupting open reading frames, whereas populations of AB5075-UW experienced a greater diversity of mutation types. IS elements are known to play an important role in *A. baumannii* mutagenesis (63). The strain-dependent historical contingencies on insertion sequence mobilization, repertoire, and insertion site could have lasting impacts on the future stability of these mutations and, therefore, the stability of resistance stemming from IS mediated mechanisms (63, 64). Further, resistance-associated mutations fixed in populations of AB5075-UW, but no mutations fixed in populations of 17978UN, indicating coexisting lineages. The maintained diversity in 17978UN lineages may be due to relatively greater mutation availability causing the coincident rise of multiple adapted lineages. When multiple mutations in the same gene collectively reach fixation, this phenomenon is known as a soft sweep (65, 66). At the population level, the diversity maintained in a population undergoing selection from antibiotics is an important indicator of future paths available for additional adaptation in new environments. In this regard, populations of 17978UN may be more evolvable when challenged with a new stress.

It would be ideal to be able to predict evolutionary trajectories of AMR from patterns of transient response to an antibiotic. While we do not see clear signatures that may enable predictions of drug adaptation at the gene level (45), our work shows that strain background greatly influences transcriptomic response to antibiotics. The large transcriptomic differences between strains in the absence of drug, which in turn influenced their responses to TGC (Fig. 1), was remarkable and could be used to map functional contingencies between strains. Many more transcriptomes of additional strains would be needed to evaluate the power of transcriptomes as markers of evolutionary history and to anticipate drug responses. Nonetheless, our studies suggest the existence of a conserved transcriptional state under TGC stress that different unexposed cell-states migrate toward. The conserved transcriptomes of evolved populations harboring several new AMR mutations further support a homogeneous state associated with drug adaptation (Fig. 5). We noted that the evolved transcriptomes more closely represent the untreated ancestral state, suggesting that the evolved populations are, essentially, not feeling the pressure of the antibiotic, likely due to the adaptations in drug efflux decreasing the intracellular concentrations of TGC (Fig. 4). It would be valuable to test if the collapse of transcriptional heterogeneity upon transient antibiotic pressure as well as prolonged adaptation is a general pattern or is contingent on the class or concentration of antibiotic applied. A lack of transcriptional response to antibiotics in clinically adapted samples could be used as another biomarker upon which to predict the extent of genetic resistance adaptation.

In a world shifting toward personalized medicine, we should also shift treatment of infections to be tailored to the specific strain or sequence type causing the infection. This work supports the idea that initial responses as well as evolutionary trajectories to antibiotic stress can be highly strain-dependent, and therefore, treatments should be as well. If we are better able to predict the impact of historical contingencies in the transcriptional response and in fitness tradeoffs associated with AMR evolution, we will be one step closer to personalized treatment of MDR infections.

## METHODS

### Bacterial strains and growth conditions

Laboratory reference strain ATCC 17978UN (38, 39) and clinical reference strain AB5075-UW (40, 41) were used throughout the study and grown under the same conditions. Unless otherwise stated, bacterial cultures were grown in 5ml M9+ media, as previously described (47). Briefly, M9+ media is a salts-buffered media with glucose (11.1mM) as the primary carbon source. It contains 0.1 mM CaCl_2_, 1 mM MgSO_4_, 42.2 mM Na_2_HPO_4_, 22 mM KH2PO_4_, 21.7 mM NaCl, 18.7 mM NH_4_Cl and is supplemented with 20 mL/L MEM essential amino acids (Gibco 11130051), 10 mL/L MEM nonessential amino acids (Gibco 11140050), and 1 mL each of trace mineral solutions A, B, and C (Corning 25021–3 Cl). Cultures were incubated at 37°C with shaking or on a roller-drum at approximately 250rpm.

### Genomic comparison of the two strains

Genomes were annotated with bakta (v1.6.1, database v4.0) (67) and input to Panaroo (v1.3) in the sensitive clean mode to obtain a gene presence/absence list that included plasmid encoded genes (68). We assessed pairwise nucleotide similarity of the two strains with pyANI (v0.2.12; (69). Multilocus sequence types were confirmed using mlst (v2.11; https://github.com/tseemann/mlst) (70). Elements associated with resistance, including known single nucleotide polymorphisms (SNPs), were detected with AMRfinderPlus (version 3.11.26 with database version 2023-11-15) (71). Two-proportion z-tests on two-tailed hypotheses were used to determine if gene content was significantly different in the two strains, accounting for the difference in genome size (Table S1). The annotated reference genomes used in this study can be found in the NCBI BioProject ID PRJNA1214285.

### Measuring resistance

Broad resistance profiles of the two strains were determined using Sensitire plates (ThermoFisher, GN3F) following the manufactures protocols. Minimum inhibitory concentration (MIC) assays were performed to measure susceptibility levels more accurately to Tigecycline (TGC, Sigma 220620-09-7) following modified CLSI methods (9, 72). More detailed methods can be found in the supplement.

### RNA extraction and purification

Cultures for transcriptomic analysis of the ancestor strains were seeded from individual colonies in biological triplicate into 5mL M9plus media. Population cultures were started directly from the freezer stock into 5mL M9plus to avoid unnecessary outgrowths. Dilutions of the cultures into M9plus with or without TGC treatment at 0.06 µg/mL were grown to mid-late exponential phase for RNA extraction. RNA was extracted following a modified TRIzol (Invitrogen Cat. No. 15596026) protocol, purified using the Invitrogen PureLink^®^ RNA Mini Kit (Thermo Cat. No. 12183025) and treated twice with DNase. See supplemental methods for more details.

### RNA sequencing and analysis

Full details for RNA sequencing and analysis can be found in the supplemental methods. Briefly, once RNA was purified and satisfied quality control measures, we performed ribosomal RNA depletion using the RiBO-COP rRNA Depletion Kit for Gram-Negative Bacteria (Lexogen Cat. No. 126.96). Library generation was done using Lexogen RNA sequencing kits, following manufacturers protocols. Sequencing was performed in house on an Illumina NextSeq550 with the corresponding High-Output Kit v2.5 75 cycles (Illumina 20024906). Raw reads are uploaded to the NCBI BioProject ID PRJNA1214285. Reads were assessed for quality, processed, and aligned to the reference genomes using kallisto (73). This analysis provided the gene abundance metrics, measured in transcripts per million transcripts (TPM), which normalizes by both gene length and sequencing depth. Differential expression analysis was done with DESeq2 (74). Unless otherwise stated, cutoffs for significance were set to a false discovery rate adjusted p-value < 0.05 and a magnitude of log_2_FoldChange > 1. RNA sequencing performed on three biological replicates for each sample by treatment combination except for removal of two replicates due to low sequencing coverage or outlier status (17978UN population 1 untreated replicate A and AB5075-UW population 2 untreated replicate C), and averages of the replicates are presented.

### Experimental evolution with TGC

Our experimental design was adapted from previous studies (47, 51). Lineages were inoculated from overnight culture with a 1:100 dilution into fresh media with or without TGC, resulting in approx. 6.6 generations per day (75). The experiment was designed such that the populations were not mutation limited (see supplemental methods). In the TGC treated condition, the TGC level in the media was tailored to the initial susceptibility level of each strain, starting at 0.5x the ancestral MIC (subinhibitory). Every three days, the concentration of TGC in the media was doubled (Fig. S5). We propagated the lineages for 12 days, with the final TGC media concentration at 4x the ancestral MIC, which crossed the clinical breakpoint for resistance in both strains. Populations were periodically sampled and frozen in 9% DMSO at - 80°C, as well as pelleted and frozen at -20°C for DNA extraction.

### Whole-population, whole-genome sequencing, and analysis

DNA was extracted from frozen cell pellets using the DNeasy blood and tissue kit for the QIAcube (Qiagen, Hilden, Germany) with a 10-minute elution into nuclease-free water. Libraries were prepared in-house as previously described (76) or using the plexWell^TM^ kit following manufacturer’s directions (SeqWell PW096). Libraries were sequenced using an Illumina NextSeq550 sequencer with a 300 cycle mid-output kit (Illumina 20024905). Raw reads are uploaded to the NCBI BioProject ID PRJNA1214285. Reads were demultiplexed, trimmed, and quality checked as described in the supplemental methods for the RNAseq reads. Breseq (v0.35.0) was used for read mapping and variant calling (77) with subsequent filtering following previously published rationale (51) with special attention to new junction calls. Filtering, consolidating for allele frequencies, and plotting were done in RStudio (R v4.2.1) with the packages ggplot2 (v3.4.2; https://CRAN.R-project.org/package=ggplot2) and tidyr (v1.3.0; https://CRAN.R-project.org/package=tidyr).

### Growth curves as measure of fitness

We use bacterial growth curves to measure absolute fitness of ancestral clonal samples as well as to measure aggregate absolute fitness of evolved populations (78). Growth curves were seeded to mimic the transfers of the evolution experiment. A large sample of the frozen population (or ancestor) stock was inoculated, in biological triplicate, into M9plus media and grown for 24h. Overnight cultures were then diluted 1:100 into fresh media and OD600 measured every 10 minutes for 24h. Fitness was measured, in technical triplicate, in plain M9plus as well as in M9plus containing 0.06 µg/mL TGC. We use area under the curve (AUC) to measure fitness of populations normalized by the AUC of their respective ancestor (AUC_evolved population_/AUC_ancestor average_). Analyses were done in RStudio (R v4.2.1) utilizing previously published pipelines (79) (https://github.com/mjfritz/Growth_Curves_in_R).

### Measurements of efflux expression and activity

Efflux activity of late exponential phase cultures was assessed using the ethidium bromide efflux activity assay. Fluorescence intensity was measured at three minutes post addition of ethidium bromide to washed cells with an excitation of 530 nm and emission of 600 nm. Quantitative reverse transcriptive PCR was used to measure expression of efflux pump components. RNA extraction was done as previously described (supplemental methods). The Power SYBR Green RNA-to-Ct 1-Step Kit (Applied Biosystems) was used for cDNA synthesis and qPCR with custom primers (Table S2). Data were analyzed for ΔCt and ΔΔCt, normalized by rpoB and ancestor, respectively, within strain background. Expression values are presented as 2^− normalized expression^ such that higher values indicate increased expression. We used the main efflux pump gene (adeB, G, and J) as a representative for pump expression (57, 80). More information on assays of efflux activity and expression can be found in supplemental methods.

## ACKNOWLEDGEMENTS

We thank all members of the Cooper Lab as well as our collaborators in the Combating Antibiotic-Resistant Bacteria Interdisciplinary Research Units (NIH CARBIRU) for thoughtful discussions and feedback. We are especially grateful for bioinformatic guidance from Emma G. Mills and Nanami Kubota. We thank Michelle R. Scribner for direction on the initial project design and assistance in uploading data to the NCBI BioProject. This work was supported by NIH U19AI158076 and, in part, by PA CURES Grant #4100085725. ABR is also supported by NIH 1F31AI172279-01A1 as well as a Catalyst Award from the Pittsburgh Center for Evolutionary Biology and Medicine (CEBaM). ASL is currently supported by the Ministry of Science, Innovation and Universities through a Ramon y Cajal fellowship (RYC2022-037765-I).

## SUPPLEMENTAL ANIMATION

**Animation S1:**
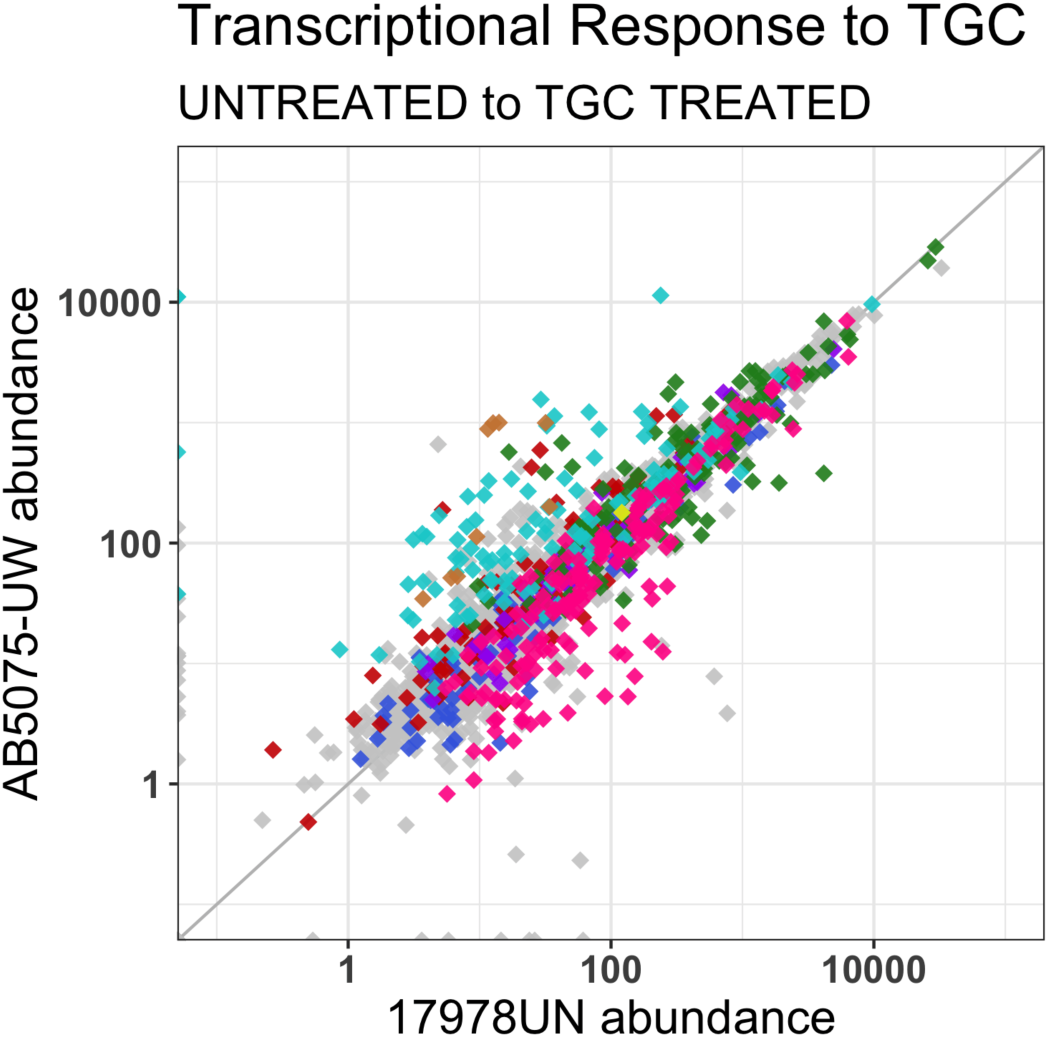
Tigecycline treatment induces a shift toward a more conserved transcriptional response. Animation of main text figure 1, showing movement of genes from the untreated state to the TGC treated state. See figure 1 for color key and legend.

## SUPPLEMENTAL FIGURES

**Fig. S1:**
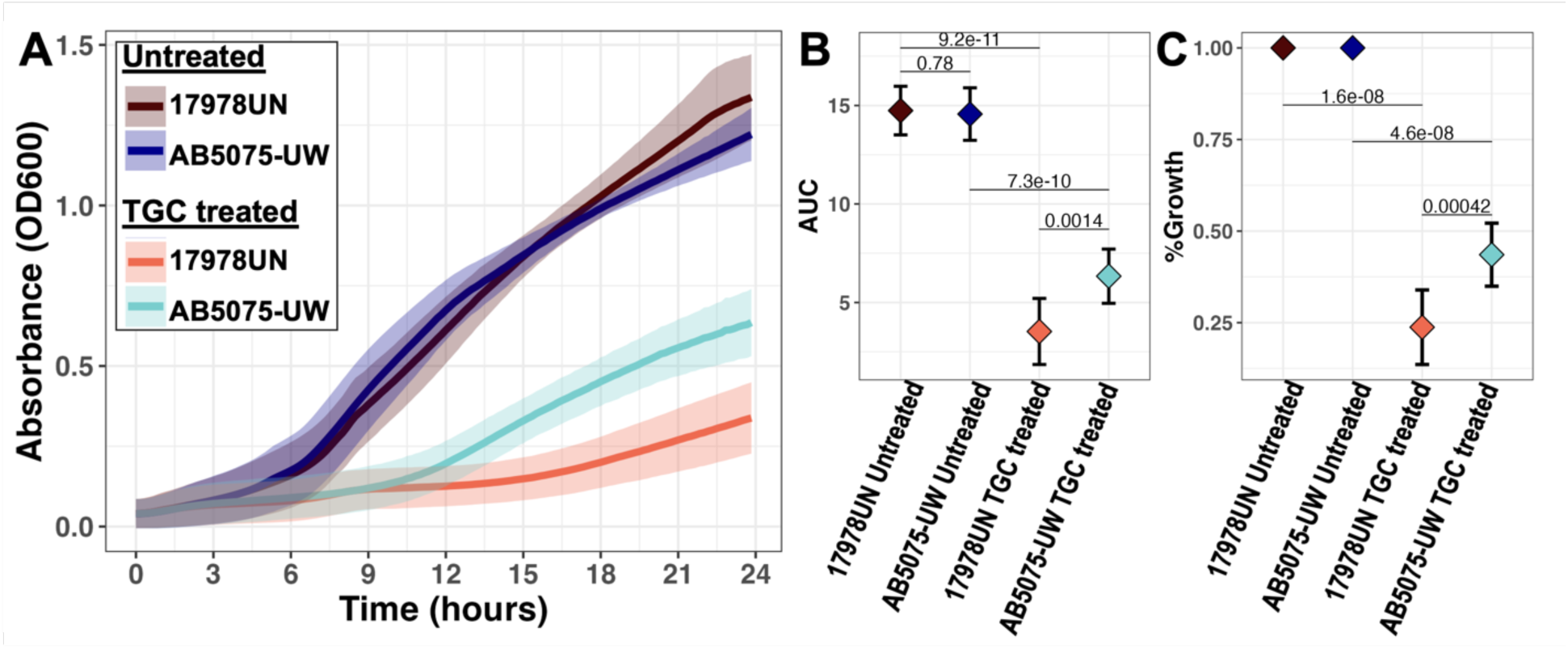
Ancestral growth in treatment conditions. Growth was measured in minimal M9plus media (Untreated) or M9plus with addition of 0.06 µg/mL TGC (TGC treated) with three biological replicates (batches), each with three technical replicates. (*A*) Growth curves for ancestors grown in different conditions. Solid line represents mean, and ribbons depict standard deviation. (*B*) Area under the curve (AUC) quantifications of curves in A (used to normalize evolved population growth in Fig. 2). (*C*) Treatment effect on ancestral growth. Y-axis shows the percent growth defect caused by TGC treatment on each strain, normalized by within-batch growth in plain media. For B and C, mean and standard deviation depicted by point and error bars, p-values calculated from pairwise t-test.

**Fig. S2:**
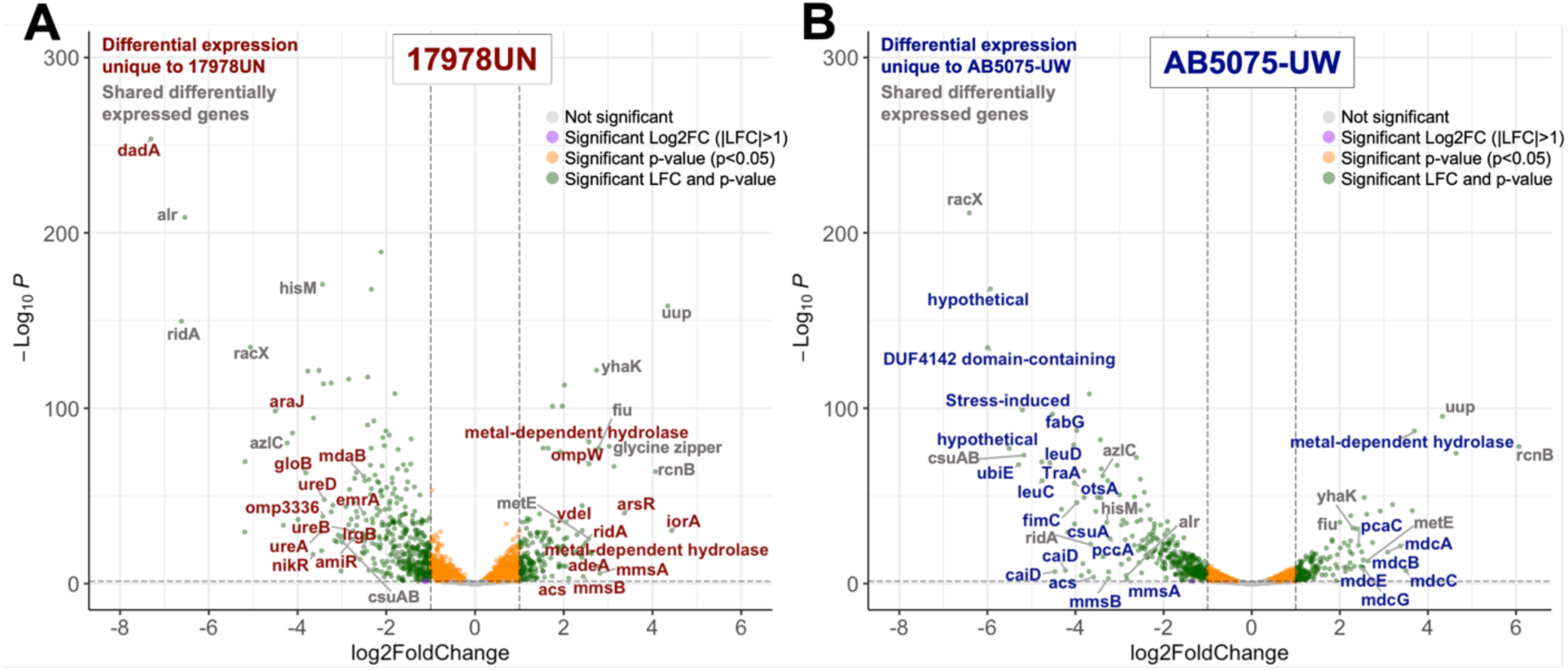
Transcriptional response to TGC is strain dependent. Volcano plots of differential expression in 17978UN (*A*) and AB5075-UW (*B*). Points are colored based on significance status and fold change; Labels are colored based on if the gene is significantly differentially expressed in only 17978UN (red), in only AB5075-UW (blue), or significant in both strains (gray). Labels were manually curated to label the strongest hits or hits that showcase differences between the strains. RNAseq was done in triplicate for each strain x treatment. Genes were included as significant in the counts/plots if they had an absolute value of log 2-fold change > 1 and a p-value ≤ 0.05.

**Fig. S3:**
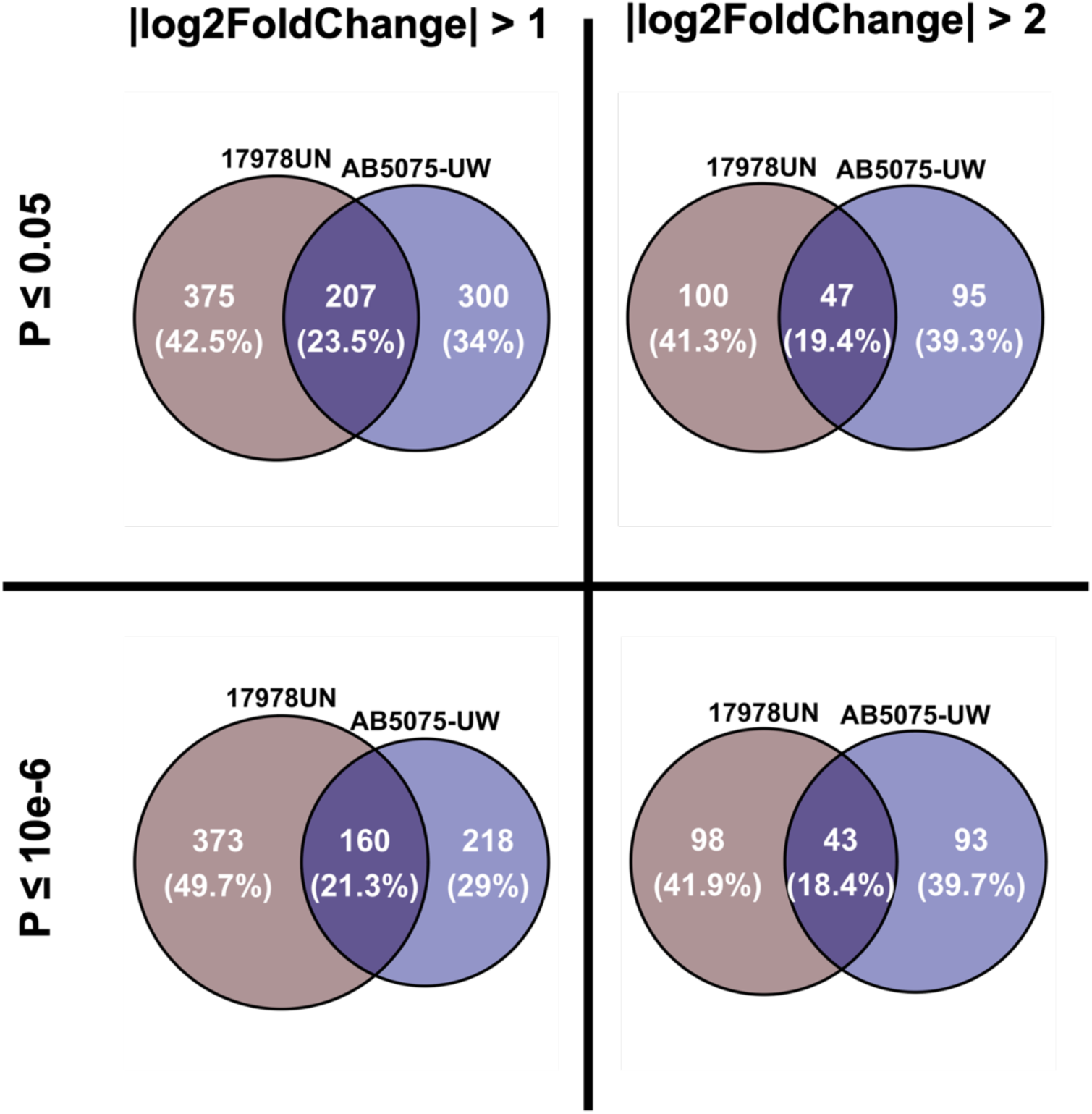
Significance cut-off criteria does not affect amount of differential expression overlap between strains. All differentially expressed genes (both upregulated and downregulated) in response to TGC treatment were summed. Varying cut-offs for calling significance in p-value and in log 2-fold change were tested to determine if the patterns of differential expression overlap between strains would differ. The overlap in differentially expressed genes between strains slightly decreases as increasingly strict significance criteria are applied.

**Fig. S4:**
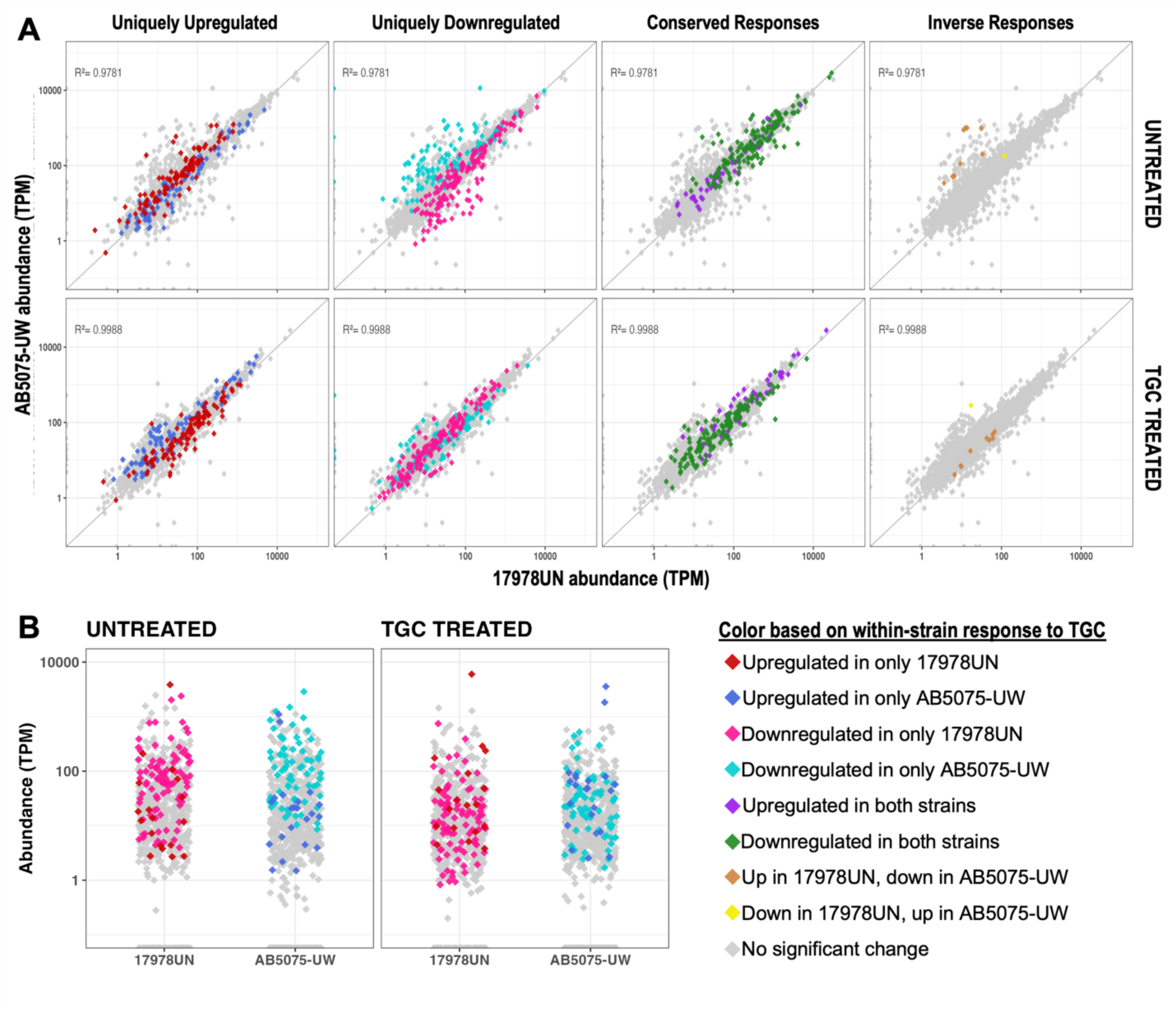
Strain-dependent transcriptional response to TGC results in a shift toward a more conserved transcriptional state. (*A*) Specific directional gene responses with genes colored based on how that gene responds within-strain upon TGC pressure. The top panels show gene abundances in untreated media and bottom panels show gene abundance in TGC treatment (0.06 µg/mL). Only genes falling under the bin of directional response noted above the plots are colored. All other genes, even those with significant TGC responses, are gray in these plots. (*B*) Abundance plots of the accessory genes, using the same colors as in A and in main text figure 1. See figure 1 legend for more details.

**Fig. S5:**
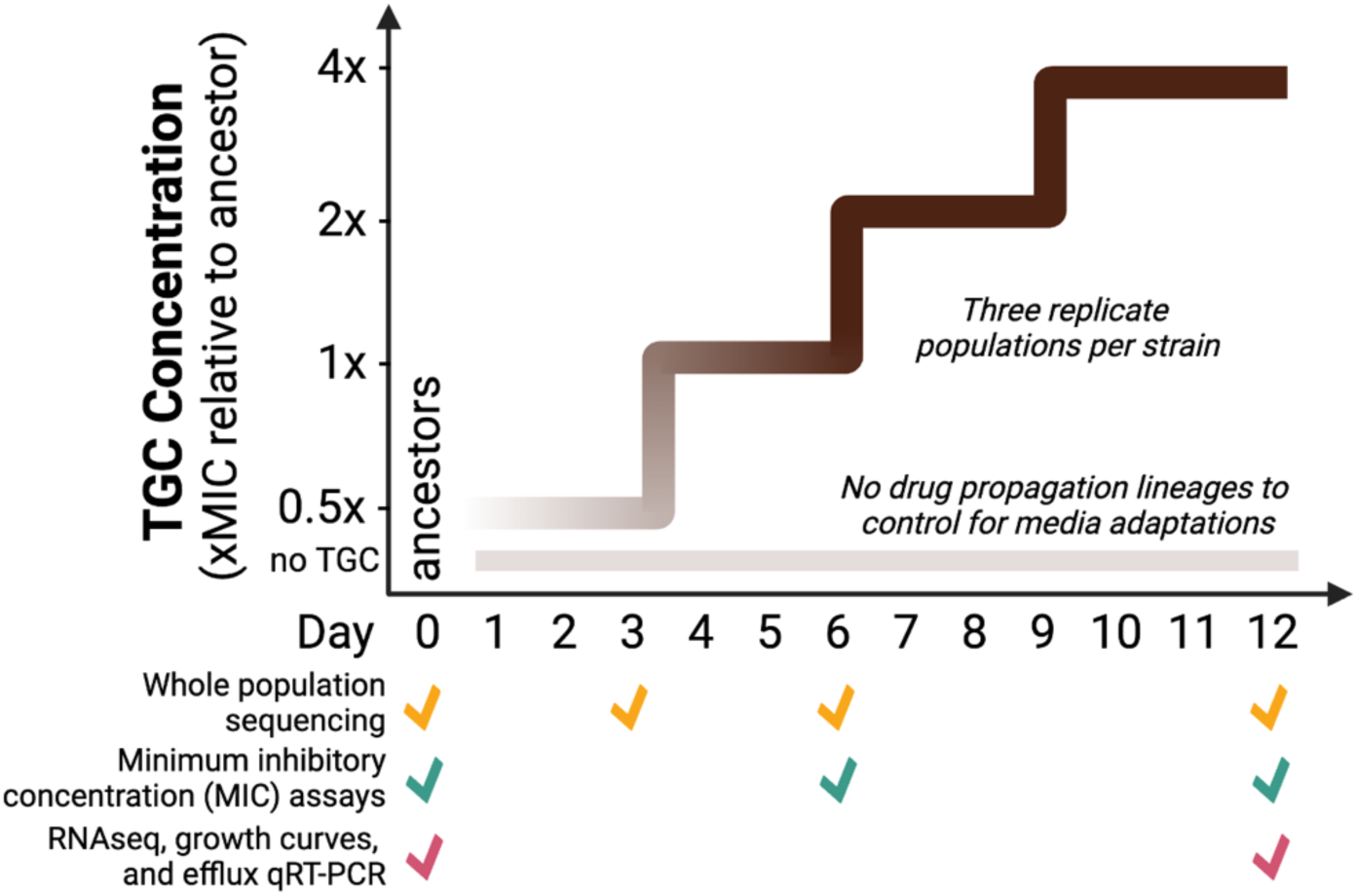
Experimental design and post-experiment assay timings. Schematic for the design of the experimental evolution. For in-depth design, see the methods section. Bottom panels highlight when various phenotype/genotype assays were performed. Dark steps indicate TGC concentration doubling every three days of the experiment. Faint bar on bottom represents lineages propagated in M9plus media lacking antibiotic to assess media adaptations.

**Fig. S6:**
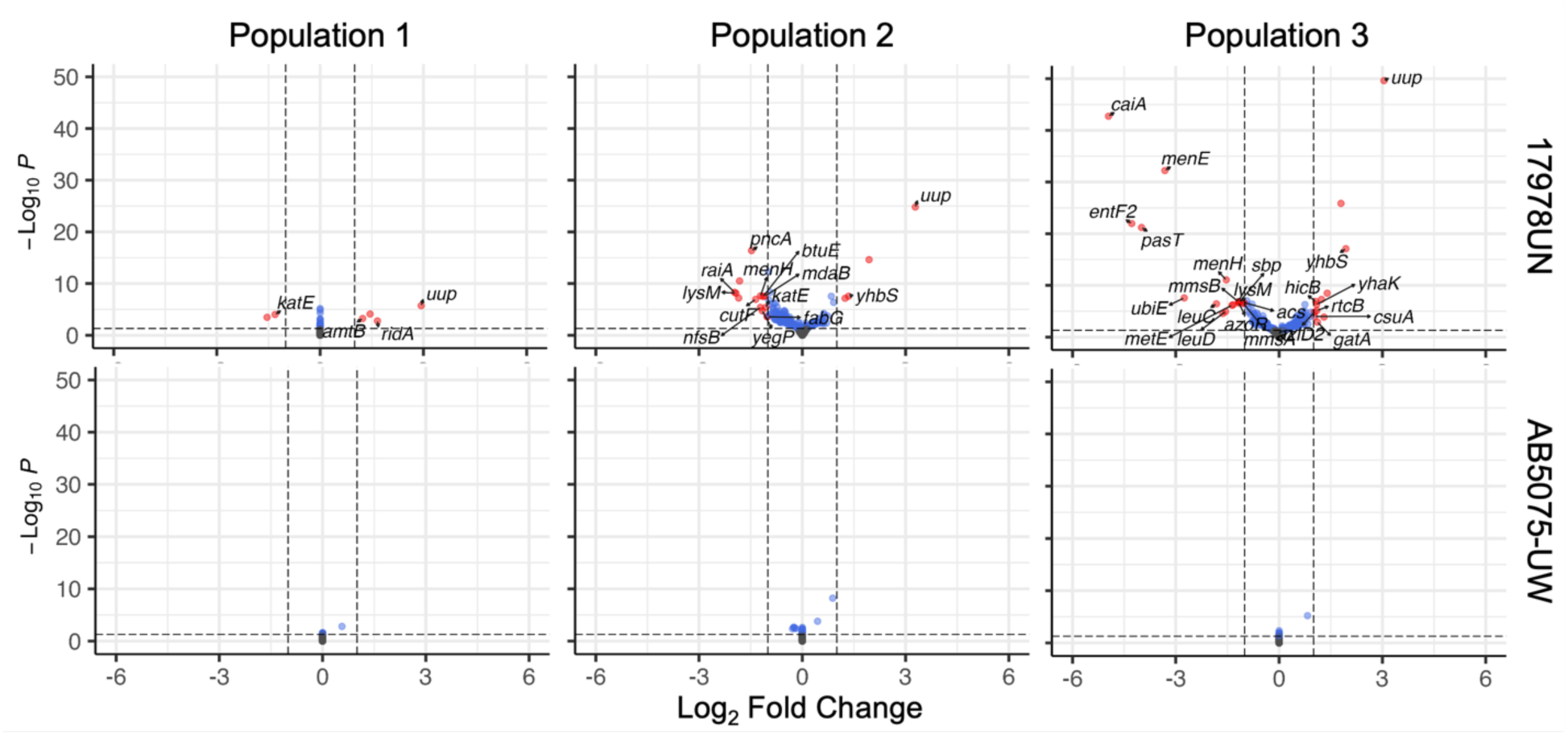
Transcriptional response to TGC in evolved populations is minimal, especially in lineages of AB5075-UW. A handful of genes significantly respond to TGC treatment (0.06 µg/mL), compared to growth in untreated media, in evolved populations of 17978UN (*top*), but no genes significantly respond to TGC in any evolved population of AB5075-UW (*bottom*). All samples are from day 12 of the TGC evolution experiment. Data is from averages of three biological replicates for all sample x treatment combinations except for removal of two replicates due to low sequencing coverage or outlier status (17978UN population 1 untreated replicate A and AB5075-UW population 2 untreated replicate C). Differential expression significance cutoffs: |LFC|>1 and p-value ≤ 0.05.

## SUPPLEMENTAL TABLES

Tables S1-S4 can also be found at https://github.com/vscooper/tigecycline

**Table S1:** Genomic comparisons between strains. Summary of strain similarities and differences in various genomic categorizations, including gene count in clusters of orthologous group (COG) categories, total genes assigned COG categories, AMR finder found resistance elements, and annotated insertion sequences. A proportional z-test was performed for all appropriate comparisons with the relevant gene count as the proportion denominator.

|  |  | AB5075-UW | 17978UN | p-value (proportional z-test) |
| --- | --- | --- | --- | --- |
| COG Categories | Cell cycle control, cell division, chromosome partitioning | 34 | 37 | 1 |
|  | Cell motility | 40 | 44 | 1 |
|  | Cell wall, membrane and envelope biogenesis | 170 | 188 | 0.957 |
|  | Cytoskeleton | 2 | 2 | 1 |
|  | Defense mechanisms | 61 | 75 | 0.547 |
|  | Extracellular structures | 36 | 39 | 1 |
|  | Intracellular trafficking, secretion, and vesicular transport | 58 | 60 | 0.832 |
|  | Posttranslational modification, protein turnover, chaperones | 115 | 128 | 0.94 |
|  | Signal transduction mechanisms | 108 | 122 | 0.855 |
|  | Replication, recombination and repair | 87 | 104 | 0.583 |
|  | RNA processing and modification | 1 | 2 | 1 |
|  | Transcription | 251 | 287 | 0.619 |
|  | Translation, ribosomal structure and biogenesis | 217 | 224 | 0.561 |
|  | Amino acid transport and metabolism | 276 | 278 | 0.33 |
|  | Carbohydrate transport and metabolism | 126 | 126 | 0.508 |
|  | Coenzyme transport and metabolism | 155 | 170 | 1 |
|  | Energy production and conversion | 164 | 170 | 0.654 |
|  | Inorganic ion transport and metabolism | 165 | 191 | 0.619 |
|  | Lipid transport and metabolism | 197 | 219 | 0.902 |
|  | Nucleotide transport and metabolism | 84 | 84 | 0.61 |
|  | Secondary metabolites biosynthesis, transport and catabolism | 55 | 68 | 0.553 |
|  | Function unknown | 113 | 130 | 0.736 |
|  | General function prediction only | 210 | 232 | 0.951 |
|  | Total number of genes with COG assignments | 2725 | 2980 | **1.55E-12** |
|  | AMRfinder Plus found elements | 19 | 6 | **0.017** |
|  | Annotated IS elements | 14 | 14 | 1 |

**Table S2:**
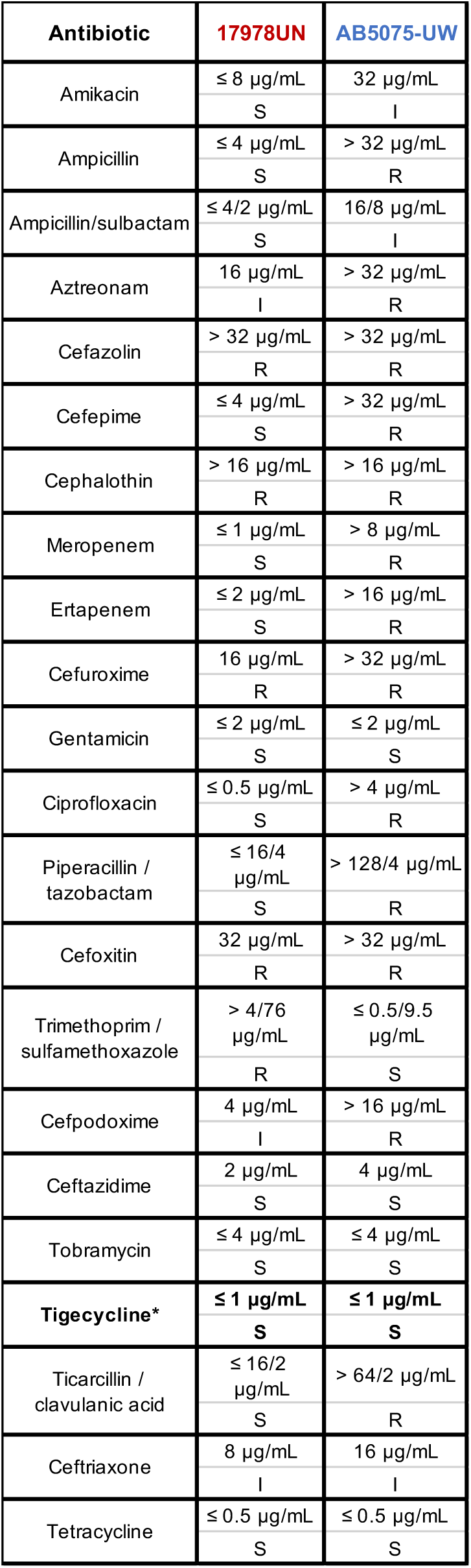
Resistance profiles for the laboratory reference strain, 17978UN, and the multi-drug resistant clinical reference strain, AB5075-UW. Susceptibility/resistance state was measured in duplicate for each strain to a variety of Gram-negative acting antibiotics via Sensititre plates #GN3F (ThermoFisher). Sensitive (S), intermediate (I), and Resistant (R) classifications were assigned using CLSI standards for Gram-negative bacteria. *Tigecycline was chosen for further use in this study and broth-dilution MICs were determined to differ 1-fold between the strains.

**Table S3:** Conservation of efflux proteins between 17978UN and AB5075-UW. Pairwise identity of amino acid sequences. * *adeC* is only encoded by AB5075-UW, but it is suggested that *adeK* can assume the function of *adeC* in 17978UN (Leus et. al., 2018). Importantly, all sites that will be mutated upon TGC adaptation are initially identical between the strains.

| Protein | Length | Amino Acid<br>Pairwise<br>Identity (%) |
| --- | --- | --- |
| AdeS | 362 | 97.2 |
| AdeR | 248 | 100 |
| AdeA | 397 | 98.7 |
| AdeB | 1037 | 99.5 |
| AdeC | 466 | * |
| AdeN | 218 | 100 |
| AdeI | 417 | 100 |
| AdeJ | 1059 | 99.9 |
| AdeK | 485 | 100 |
| AdeL | 344 | 99.7 |
| AdeF | 407 | 99.5 |
| AdeG | 1060 | 99.9 |
| AdeH | 483 | 99.8 |

**Table S4:**
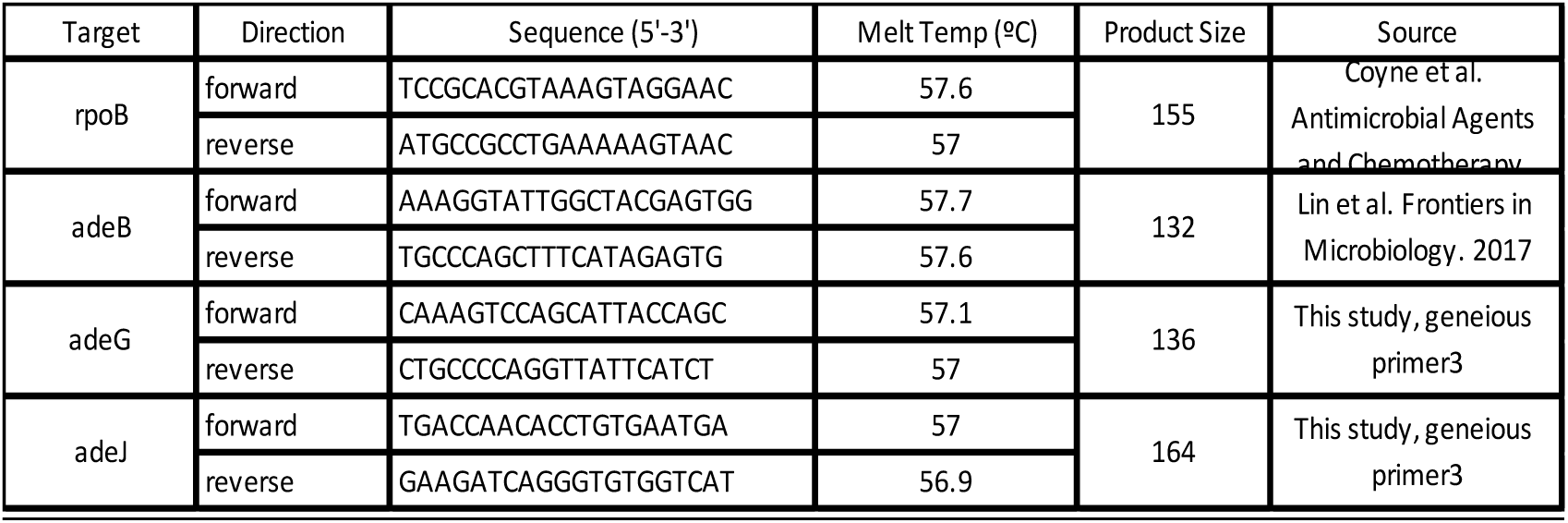
List of primers used for Q-RT-PCR. Quantative, reverse-transcriptase PCR was performed on the inner membrane pump of the three RND efflux pumps as documented in the methods using primers with the characteristics listed here. The housekeeping gene *rpoB* was used as a control.

## SUPPLEMENTAL DATA LEGENDS

*There are three submissions for supplemental data, each a separate file that can be accessed through https://github.com/vscooper/tigecycline*

Please contact authors for additional information about supplemental data.

**Data S1: Ancestral gene annotations and RNA sequencing results.**

**Data S2: Filtered results of whole-population, whole-genome sequencing throughout the evolution experiment.**

**Data S3: RNA sequencing results of the evolved populations.**

## SUPPLEMENTAL METHODS

Supplemental methods citations found after main text citations.

### Bacterial strains and growth conditions

Laboratory reference strain ATCC 17978UN (1, 2) and clinical reference strain AB5075-UW (3, 4) were used throughout the study and grown under the same conditions. Unless otherwise stated, bacterial cultures were grown in 5ml M9+ media, as previously described (5). Briefly, M9+ media is a salts-buffered media with glucose (11.1mM) as the primary carbon source. It contains 0.1 mM CaCl_2_, 1 mM MgSO_4_, 42.2 mM Na_2_HPO_4_, 22 mM KH2PO_4_, 21.7 mM NaCl, 18.7 mM NH_4_Cl and is supplemented with 20 mL/L MEM essential amino acids (Gibco 11130051), 10 mL/L MEM nonessential amino acids (Gibco 11140050), and 1 mL each of trace mineral solutions A, B, and C (Corning 25021–3 Cl). Cultures were incubated at 37°C with shaking or on a roller-drum at approximately 250rpm.

### Genomic comparison of the two strains

Genomes were annotated with bakta (v1.6.1, database v4.0) (6) and input to Panaroo (v1.3) in the sensitive clean mode to obtain a gene presence/absence list that included plasmid encoded genes (Data S1) (7). We assessed pairwise nucleotide similarity of the two strains with pyANI (v0.2.12; (8). Multilocus sequence types were confirmed using mlst (v2.11; https://github.com/tseemann/mlst) (9). Elements associated with resistance, including known single nucleotide polymorphisms (SNPs), were detected with AMRfinderPlus (version 3.11.26 with database version 2023-11-15) (10). Two-proportion z-tests on two-tailed hypotheses were used to determine if gene content in clusters of orthologous group (COGs) categories was significantly different in the two strains, accounting for the difference in genome size (Table S1) (11). The annotated reference genomes used in this study can be found in the NCBI BioProject ID PRJNA1214285.

### Measuring resistance

Broad resistance profiles of the two strains were determined using Sensititre plates (ThermoFisher, GN3F), commercially available panels of 23 antibiotics relevant to Gram-negative pathogens at dilutions spanning the clinical breakpoint for resistance as set by the Clinical and Laboratory Standards Institute (CLSI) (12). Broth microdilution and inoculation of the plates was performed following the manufacturer’s instructions. Sensititre plates were performed in biological triplicate for each strain.

Minimum inhibitory concentration (MIC) assays were performed to measure susceptibility levels more accurately to tigecycline (TGC, Sigma 220620-09-7). MIC assays were performed following modified CLSI methods (12). MIC results presented here are from MIC assays performed in M9plus media. Reference MIC assays were performed in the standard Mueller Hinton broth and showed little-to-no (maximum difference of 1-fold) difference to those performed in M9plus media. Biological triplicate (multiple samples from the frozen ancestors or populations) with technical triplicates for each assay were performed. We present the concentration of antibiotic that inhibits 90% growth (IC_90_), measured via optical density at 600nm, as the MIC for each antibiotic.

### RNA extraction and purification

For transcriptome analysis of the ancestor strains, cultures were seeded from individual colonies in biological triplicate into 5mL M9plus media. Population cultures were started directly by inoculating a large sampling of freezer stock into 5mL M9plus, also in triplicate. All cultures were grown over-night at 37°C and back diluted 1:100 into 25 mL pre-warmed M9plus with or without 0.06 µg/mL TGC. In this way, each biological replicate over-night culture results in a paired set of treated and untreated TGC culture for analysis, thus limiting the effect of sampling bias for the populations. Cultures were grown until reaching mid-log phase (OD600 = 0.6 +/- 0.05), at which time cultures were pelleted, resuspended in TRIzol (Invitrogen Cat. No. 15596026) and stored at -80°C for subsequent extraction and purification.

RNA was extracted following a modified protocol (13) for TRIzol extraction with purification using the Invitrogen PureLink^®^ RNA Mini Kit. Extraction included one bead beating step followed by TRIzol-chloroform extraction with ethanol precipitation. Purification was done using RNA columns from the Invitrogen PureLink system with final elution into nuclease free water. RNA samples were treated with DNase 1 off-column with additional DNase spike-in during incubation (NEB DNase and buffer Cat. No. M0303S) as well as treated with DNase 1 on-column (Qiagen Cat. No. 79254) during final purification steps on the Invitrogen columns. RNA concentration and quality were assessed using nanodrop (all 260/280 ratios > 2). Pure RNA was stored at -20°C for less than a week prior to library preparation and at -80°C for longer-term storage. Step-by-step RNA protocols are available upon request.

### RNA sequencing and analysis

RNA integrity, fragment distribution, and RIN^e^ scores were assessed on Tapestation using the High Sensitivity RNA ScreenTape, sample buffer, and ladder from Agilent Technologies (Part Numbers: 5067 -5579, -5580, -5581). All RIN^e^ scores satisfied subsequent ribo-depeletion requirements for quality (the majority of RIN^e^ scores were >8). rRNA depletion was performed using the RiBO-COP rRNA Depletion Kit for Gram-Negative Bacteria (Lexogen Cat. No. 126.96) and directly following the manufacturer’s protocols with an initial RNA input of approximately 200ng. RNA libraries including ancestral strain RNAs were prepared using the CORALL Total RNA-Seq Library Prep Kit (Lexogen Cat. No. 095) while the evolved populations were sequenced with the RNA-Seq V2 Library Prep Kit with UDIs (Lexogen Cat. No. 171.96). Both library preparations included the PCR add-on step to determine ideal cycle number (11 cycles) for library amplification (Lexogen Cat. No. 020). The uniquely indexed samples for each library were pooled based on fragment size and concentration and the pool was diluted to 4nM for sequencing. Each library was denatured and loaded at a final concentration of 2pM with a 5% phiX spike-in. Sequencing was performed on the Illumina NextSeq550 with the corresponding High-Output Kit v2.5 75 cycles (Illumina 20024906). Raw reads are uploaded to the NCBI BioProject ID PRJNA1214285.

Reads were demultiplexed using bcl2fastq (v2.17.1) and trimmed with trimmomatic (v0.36, custom criteria as follows: LEADING:3 TRAILING:3 SLIDINGWINDOW:4:15 MINLEN:36). Read quality was confirmed with fastQC (v0.11.5) and kallisto (v0.48.0, flags as follows: --single -l 315 -s 26) was used for pseudoalignment with custom made index files from the concatenated bakta annotated fasta files previously described for each reference strain (14). Sequencing coverage was satisfactory (>10^7^ reads/sample, resulting in >200x coverage) for all samples apart from one replicate of AB5075-UW evolved population 2 in the untreated condition, which had unusually low coverage and was removed from further analysis. We also removed one replicate of 17978UN evolved population 1 in the untreated condition. DESeq2 (v1.38.3; (15)) was used in RStudio (R v4.2.1) for differential expression analysis supplemented with EnhancedVolcano (v1.16.0) for plotting. Transcript abundance is reported as transcripts per million transcripts (TPM) which normalizes by both gene length and sequencing depth. TGC imposed differential expression analyses were performed with guidance from established tutorials (15, 16). In the results function for analysis of the DESeq Data Set, we specified to turn off independent filtering and cooksCutoff to keep outliers and calls with low coverage in the analyses, then the apeglm method was applied for log fold change shrinkage (17). Unless otherwise stated, cutoffs for significance were set to a false discovery rate adjusted p-value < 0.05, with the Benjamini and Hochberg correction for multiple comparisons, and a magnitude of log_2_FoldChange > 1.

### Experimental evolution with TGC

Our experimental design was adapted from previous studies (5, 18). Both strains were resurrected from the freezer via streaking on LB agar plates. A single colony of each strain was used to inoculate a 5mL overnight culture in M9+. After approximately 18 hours, the saturated culture was used to inoculate multiple independent lineages for the evolution experiment. We propagated five replicate lineages of each strain in each condition, either with or without TGC. Lineages were inoculated from overnight culture (either the starting overnight, or the prior day) with a 1:100 dilution into fresh media with or without TGC. 1:100 dilutions from saturated overnight cultures results in approximately 6.6 generations per day (19). The large population size combined with the 1% bottleneck, mutation rate, and doubling time of *A. baumannii* provides a large mutation supply, with a probability >1 that each site can see a mutation (19).

Lineages were propagated in the presence and absence of TGC, resulting in two conditions. The no drug lineages were used to control for adaptations to the media and experimental conditions. In the TGC treated condition, the TGC level in the media was tailored to the initial susceptibility level of each strain, starting at 0.5x the ancestral MIC (subinhibitory). Every three days, the concentration of TGC in the media was doubled (Fig. S5). We propagated the lineages for 12 days, with the final TGC media concentration at 4x the ancestral MIC, which crossed the clinical breakpoint for resistance in both strains. On days 0, 1, 3, 4, 6, 7, 9, 10, 12, corresponding to days before and after antibiotic level increases, the populations were stocked for later assays and sequencing (frozen in 9% DMSO at -80°C, and a cell pellet from 1mL of culture was frozen at -20°C).

### Whole-population, whole-genome sequencing, and analysis

DNA was extracted from frozen cell pellets using the DNeasy blood and tissue kit for the QIAcube (Qiagen, Hilden, Germany) with a 10-minute elution into nuclease-free water. Libraries were prepared in-house as previously described (20, 21) or using the plexWell^TM^ kit following manufacturer’s directions (SeqWell PW096). Libraries were sequenced using an Illumina NextSeq550 sequencer with a 300 cycle mid-output kit (Illumina 20024905). Reads were demultiplexed, trimmed, and quality checked as described in the supplemental methods for the RNAseq reads. Raw reads are uploaded to NCBI BioProject ID PRJNA1214285. Breseq (v0.35.0) was used for read mapping and variant calling (22) with subsequent filtering following previously published rationale (18) with special attention to new junction calls. Filtering, consolidating for allele frequencies, and plotting were done in RStudio (R v4.2.1) with the packages ggplot2 (v3.4.2; https://CRAN.R-project.org/package=ggplot2) and tidyr (v1.3.0; https://CRAN.R-project.org/package=tidyr).

### Growth curves as measure of fitness

We use bacterial growth curves to measure absolute fitness of ancestral clonal samples as well as to measure aggregate absolute fitness of evolved populations (23). Growth curves were seeded to mimic the transfers of the evolution experiment. A large sample of the frozen population (or ancestor) stock was inoculated, in biological triplicate, into M9plus media and grown for 24h. Overnight cultures were then diluted 1:100 into fresh media and OD600 measured every 10 minutes for 24h. Fitness was measured, in technical triplicate, in plain M9plus as well as in M9plus containing 0.06 µg/mL TGC. We use area under the curve (AUC) to measure fitness of populations normalized by the AUC of their respective ancestor (AUC_evolved population_/AUC_ancestor average_). Analyses were done in RStudio (R v4.2.1) utilizing previously published pipelines (24) (https://github.com/mjfritz/Growth_Curves_in_R).

### Quantitative reverse transcriptive PCR

To measure evolved efflux pump expression, we extracted RNA as described above for ancestors and day-12 TGC evolved populations grown in the presence (0.06 µg/mL TGC, ancestral cultures only) or absence of TGC (for ancestral and evolved populations). 50ng of purified RNA was added per reaction for Q-RT-PCR according to manufacturer’s instructions for the Power SYBR Green RNA-to-Ct 1-Step Kit (Applied Biosystems). Reactions were performed in biological and technical triplicate and run on a QuantStudio3 thermocycler using the default threshold detection. Reactions lacking reverse transcriptase mix were included as controls for DNA contamination. Data were analyzed for ΔCt and ΔΔCt, normalized by *rpoB* and ancestor, respectively, within strain background. Expression values are presented as 2^−normalized expression^ such that higher values indicate increased expression. We used the main efflux pump gene (*adeB, G, and J)* as a representative for pump expression (25, 26). Primers were designed with Geneious to amplify approximately 150 base pair regions and are listed in Table S4.

### Ethidium bromide efflux activity assay

Efflux activity was measured with the ethidium bromide uptake assay loosely based on previously published methods (27). Ancestral cultures were grown overnight in M9plus, either with or without addition of 0.06 µg/mL TGC, but not until saturation, and 500 µL was pelleted, washed twice, and resuspended with PBS. Efflux activity of day-12 TGC-evolved populations was measured only in untreated M9plus. A black flat-bottom 96-well plate (Corning ref. 3925) was seeded with 90 µL of resuspended cells and 10 µL of 10 µg/mL ethidium bromide (Invitrogen by Thermo Fisher Scientific ref. 15585-011). Fluorescence intensity was measured at three minutes post addition of ethidium bromide with an excitation of 530 nm and emission of 600 nm. High efflux activity would result in a low fluorescence intensity as less ethidium bromide is left within the cells binding to DNA. We present efflux activity as 1/RFU such that higher values indicate more efflux. Efflux activity of the day-12 TGC-evolved populations when grown in the untreated state relative to respective ancestor was calculated as 1/(RFU_query_/RFU_untreated ancestor_). Efflux activity was measured in biological triplicate with at least three technical replicates each.

## Main Text Citations

1. B. Aslam, et al., Antibiotic resistance: a rundown of a global crisis. Infect. Drug Resist. 11, 1645–1658 (2018).

2. M. Lässig, V. Mustonen, A. M. Walczak, Predicting evolution. Nat. Ecol. Evol. 1, 77 (2017).

3. T. Gabaldón, Nothing makes sense in drug resistance except in the light of evolution. Curr. Opin. Microbiol. 75, 102350 (2023).

4. A. C. Palmer, R. Kishony, Understanding, predicting and manipulating the genotypic evolution of antibiotic resistance. Nat. Rev. Genet. 14, 243–248 (2013).

5. M. Travisano, J. A. Mongold, A. F. Bennett, R. E. Lenski, Experimental tests of the roles of adaptation, chance, and history in evolution. Science 267, 87–90 (1995).

6. J. Plucain, et al., Contrasting effects of historical contingency on phenotypic and genomic trajectories during a two-step evolution experiment with bacteria. BMC Evol. Biol. 16, 86 (2016).

7. R. C. MacLean, A. San Millan, The evolution of antibiotic resistance. Science 365, 1082–1083 (2019).

8. Z. D. Blount, R. E. Lenski, J. B. Losos, Contingency and determinism in evolution: Replaying life’s tape. Science 362 (2018).

9. A. Santos-Lopez, et al., The roles of history, chance, and natural selection in the evolution of antibiotic resistance. eLife 10 (2021).

10. L. Garoff, et al., Population Bottlenecks Strongly Influence the Evolutionary Trajectory to Fluoroquinolone Resistance in Escherichia coli. Mol. Biol. Evol. 37, 1637–1646 (2020).

11. A. Wong, Epistasis and the evolution of antimicrobial resistance. Front. Microbiol. 8, 246 (2017).

12. R. C. MacLean, Predicting epistasis: an experimental test of metabolic control theory with bacterial transcription and translation. J. Evol. Biol. 23, 488–493 (2010).

13. Z. D. Blount, A case study in evolutionary contingency. Stud. Hist. Philos. Biol. Biomed. Sci. 58, 82–92 (2016).

14. G. J. Vermeij, Historical contingency and the purported uniqueness of evolutionary innovations. Proc Natl Acad Sci USA 103, 1804–1809 (2006).

15. Y. Wang, et al., Benefit of transferred mutations is better predicted by the fitness of recipients than by their ecological or genetic relatedness. Proc Natl Acad Sci USA 113, 5047–5052 (2016).

16. S. Dionisi, J. S. Huisman, Digest: The importance of genetic background in bacterial evolution. Evolution 78, 593–594 (2024).

17. N. Filipow, S. Mallon, S. Shewaramani, R. Kassen, A. Wong, The impact of genetic background during laboratory evolution of Pseudomonas aeruginosa in a cystic fibrosis-like environment. Evolution 78, 566–578 (2024).

18. S. Cisneros-Mayoral, L. Graña-Miraglia, D. Pérez-Morales, R. Peña-Miller, A. Fuentes-Hernández, Evolutionary history and strength of selection determine the rate of antibiotic resistance adaptation. Mol. Biol. Evol. 39 (2022).

19. K. J. Card, T. LaBar, J. B. Gomez, R. E. Lenski, Historical contingency in the evolution of antibiotic resistance after decades of relaxed selection. PLoS Biol. 17, e3000397 (2019).

20. T. N. Batarseh, S. N. Batarseh, A. Rodríguez-Verdugo, B. S. Gaut, Phenotypic and Genotypic Adaptation of Escherichia coli to Thermal Stress is Contingent on Genetic Background. Mol. Biol. Evol. 40 (2023).

21. K. M. Flynn, T. F. Cooper, F. B.-G. Moore, V. S. Cooper, The environment affects epistatic interactions to alter the topology of an empirical fitness landscape. PLoS Genet. 9, e1003426 (2013).

22. F. Baquero, et al., Evolutionary pathways and trajectories in antibiotic resistance. Clin. Microbiol. Rev. 34, e0005019 (2021).

23. C.-R. Lee, et al., Biology of Acinetobacter baumannii: Pathogenesis, Antibiotic Resistance Mechanisms, and Prospective Treatment Options. Front. Cell. Infect. Microbiol. 7, 55 (2017).

24. M. Touchon, et al., The genomic diversification of the whole Acinetobacter genus: origins, mechanisms, and consequences. Genome Biol. Evol. 6, 2866–2882 (2014).

25. A. Iovleva, et al., Carbapenem-Resistant Acinetobacter baumannii in U.S. Hospitals: Diversification of Circulating Lineages and Antimicrobial Resistance. MBio 13, e0275921 (2022).

26. M. J. Dorman, N. R. Thomson, “Community evolution” - laboratory strains and pedigrees in the age of genomics. *Microbiology (Reading*, Engl) 166, 233–238 (2020).

27. M. Del Mar Cendra, E. Torrents, Differential adaptability between reference strains and clinical isolates of Pseudomonas aeruginosa into the lung epithelium intracellular lifestyle. Virulence 11, 862–876 (2020).

28. G. Makke, et al., Whole-Genome-Sequence-Based Characterization of Extensively Drug-Resistant Acinetobacter baumannii Hospital Outbreak. mSphere 5 (2020).

29. C. H. Camargo, et al., Clonal spread of ArmA- and OXA-23-coproducing Acinetobacter baumannii International Clone 2 in Brazil during the first wave of the COVID-19 pandemic. J. Med. Microbiol. 71 (2022).

30. M. Marazzato, et al., Genetic Diversity of Antimicrobial Resistance and Key Virulence Features in Two Extensively Drug-Resistant Acinetobacter baumannii Isolates. Int. J. Environ. Res. Public Health 19 (2022).

31. M. R. Galac, et al., A Diverse Panel of Clinical Acinetobacter baumannii for Research and Development. Antimicrob. Agents Chemother. 64 (2020).

32. T. van Opijnen, S. Dedrick, J. Bento, Strain Dependent Genetic Networks for Antibiotic-Sensitivity in a Bacterial Pathogen with a Large Pan-Genome. PLoS Pathog. 12, e1005869 (2016).

33. L. Rizzetto, et al., Strain dependent variation of immune responses to A. fumigatus: definition of pathogenic species. PLoS ONE 8, e56651 (2013).

34. J. Davies, D. Davies, Origins and evolution of antibiotic resistance. Microbiol. Mol. Biol. Rev. 74, 417–433 (2010).

35. A. Kaul, C. Souque, M. Holland, M. Baym, Genomic resistance in historical clinical isolates increased in frequency and mobility after the age of antibiotics. BioRxiv (2025) 10.1101/2025.01.16.633422.

36. A. P. Hendry, Prediction in ecology and evolution. Bioscience (2023) 10.1093/biosci/biad083.

37. K. E. Erickson, P. B. Otoupal, A. Chatterjee, Transcriptome-Level Signatures in Gene Expression and Gene Expression Variability during Bacterial Adaptive Evolution. mSphere 2 (2017).

38. C. D. M. Wijers, et al., Identification of Two Variants of Acinetobacter baumannii Strain ATCC 17978 with Distinct Genotypes and Phenotypes. Infect. Immun. 89, e0045421 (2021).

39. P. Baumann, M. Doudoroff, R. Y. Stanier, A study of the Moraxella group. II. Oxidative-negative species (genus Acinetobacter). J. Bacteriol. 95, 1520–1541 (1968).

40. A. C. Jacobs, et al., AB5075, a Highly Virulent Isolate of Acinetobacter baumannii, as a Model Strain for the Evaluation of Pathogenesis and Antimicrobial Treatments. MBio 5, e01076–14 (2014).

41. L. A. Gallagher, et al., Resources for Genetic and Genomic Analysis of Emerging Pathogen Acinetobacter baumannii. J. Bacteriol. 197, 2027–2035 (2015).

42. N. D. Greer, Tigecycline (Tygacil): the first in the glycylcycline class of antibiotics. Proc (Bayl Univ Med Cent) 19, 155–161 (2006).

43. X. Hua, et al., Novel tigecycline resistance mechanisms in Acinetobacter baumannii mediated by mutations in adeS, rpoB and rrf. Emerg. Microbes Infect. 10, 1404–1417 (2021).

44. K. Lucaßen, et al., Prevalence of RND efflux pump regulator variants associated with tigecycline resistance in carbapenem-resistant Acinetobacter baumannii from a worldwide survey. J. Antimicrob. Chemother. 76, 1724–1730 (2021).

45. X. Hua, Q. Chen, X. Li, Y. Yu, Global transcriptional response of Acinetobacter baumannii to a subinhibitory concentration of tigecycline. Int. J. Antimicrob. Agents 44, 337–344 (2014).

46. L. Gao, X. Ma, Transcriptome Analysis of Acinetobacter baumannii in Rapid Response to Subinhibitory Concentration of Minocycline. Int. J. Environ. Res. Public Health 19 (2022).

47. A. Santos-Lopez, C. W. Marshall, M. R. Scribner, D. J. Snyder, V. S. Cooper, Evolutionary pathways to antibiotic resistance are dependent upon environmental structure and bacterial lifestyle. eLife 8 (2019).

48. T. Vogwill, M. Kojadinovic, R. C. MacLean, Epistasis between antibiotic resistance mutations and genetic background shape the fitness effect of resistance across species of Pseudomonas. Proc. Biol. Sci. 283 (2016).

49. T. Vogwill, R. C. MacLean, The genetic basis of the fitness costs of antimicrobial resistance: a meta-analysis approach. Evol. Appl. 8, 284–295 (2015).

50. K. J. Card, J. A. Jordan, R. E. Lenski, Idiosyncratic variation in the fitness costs of tetracycline-resistance mutations in Escherichia coli. Evolution 75, 1230–1238 (2021).

51. M. R. Scribner, A. Santos-Lopez, C. W. Marshall, C. Deitrick, V. S. Cooper, Parallel Evolution of Tobramycin Resistance across Species and Environments. MBio 11 (2020).

52. K. B. Harris, K. M. Flynn, V. S. Cooper, Polygenic adaptation and clonal interference enable sustained diversity in experimental Pseudomonas aeruginosa populations. Mol. Biol. Evol. (2021) 10.1093/molbev/msab248.

53. Q. Chen, et al., Decreased susceptibility to tigecycline in Acinetobacter baumannii mediated by a mutation in trm encoding SAM-dependent methyltransferase. J. Antimicrob. Chemother. 69, 72–76 (2014).

54. S. Sharma, V. Kaushik, M. Kulshrestha, V. Tiwari, Different Efflux Pump Systems in Acinetobacter baumannii and Their Role in Multidrug Resistance. Adv. Exp. Med. Biol. 1370, 155–168 (2023).

55. I. V. Leus, et al., Substrate Specificities and Efflux Efficiencies of RND Efflux Pumps of Acinetobacter baumannii. J. Bacteriol. 200 (2018).

56. S. Raza, et al., Molecular Characterization of Resistance-Nodulation-cell Division Efflux Pump Genes in Multidrug-Resistant Acinetobacter baumannii. J. Glob. Infect. Dis. 13, 177–179 (2021).

57. S. Coyne, P. Courvalin, B. Périchon, Efflux-mediated antibiotic resistance in Acinetobacter spp. Antimicrob. Agents Chemother. 55, 947–953 (2011).

58. W. Huo, et al., Immunosuppression broadens evolutionary pathways to treatment failure during *Acinetobacter baumannii* pneumonia. BioRxiv (2021) 10.1101/2021.04.07.438861.

59. N. Harmand, R. Gallet, G. Martin, T. Lenormand, Evolution of bacteria specialization along an antibiotic dose gradient. Evol. Lett. 2, 221–232 (2018).

60. L. Praski Alzrigat, D. L. Huseby, G. Brandis, D. Hughes, Resistance/fitness trade-off is a barrier to the evolution of MarR inactivation mutants in Escherichia coli. J. Antimicrob. Chemother. 76, 77–83 (2021).

61. B. R. Levin, V. Perrot, N. Walker, Compensatory mutations, antibiotic resistance and the population genetics of adaptive evolution in bacteria. Genetics 154, 985–997 (2000).

62. E. M. Darby, V. N. Bavro, S. Dunn, A. McNally, J. M. A. Blair, RND pumps across the genus Acinetobacter: AdeIJK is the universal efflux pump. Microb. Genom. 9 (2023).

63. S. Gaiarsa, et al., Can Insertion Sequences Proliferation Influence Genomic Plasticity? Comparative Analysis of Acinetobacter baumannii Sequence Type 78, a Persistent Clone in Italian Hospitals. Front. Microbiol. 10, 2080 (2019).

64. M. S. Wright, S. Mountain, K. Beeri, M. D. Adams, Assessment of Insertion Sequence Mobilization as an Adaptive Response to Oxidative Stress in Acinetobacter baumannii Using IS-seq. J. Bacteriol. 199 (2017).

65. B. H. Good, I. M. Rouzine, D. J. Balick, O. Hallatschek, M. M. Desai, Distribution of fixed beneficial mutations and the rate of adaptation in asexual populations. Proc Natl Acad Sci USA 109, 4950–4955 (2012).

66. M. M. Desai, D. S. Fisher, Beneficial mutation selection balance and the effect of linkage on positive selection. Genetics 176, 1759–1798 (2007).

67. O. Schwengers, et al., Bakta: rapid and standardized annotation of bacterial genomes via alignment-free sequence identification. Microb. Genom. 7 (2021).

68. G. Tonkin-Hill, et al., Producing polished prokaryotic pangenomes with the Panaroo pipeline. Genome Biol. 21, 180 (2020).

69. L. Pritchard, R. H. Glover, S. Humphris, J. G. Elphinstone, I. K. Toth, Genomics and taxonomy in diagnostics for food security: soft-rotting enterobacterial plant pathogens. Anal. Methods 8, 12–24 (2016).

70. K. A. Jolley, M. C. J. Maiden, BIGSdb: Scalable analysis of bacterial genome variation at the population level. BMC Bioinformatics 11, 595 (2010).

71. M. Feldgarden, et al., AMRFinderPlus and the Reference Gene Catalog facilitate examination of the genomic links among antimicrobial resistance, stress response, and virulence. Sci. Rep. 11, 12728 (2021).

72. C. CLSI, M100: Performance Standards for Antimicrobial Susceptibility Testing, 32nd Ed. (Clinical and Laboratory Standards Institute, 2022).

73. N. L. Bray, H. Pimentel, P. Melsted, L. Pachter, Near-optimal probabilistic RNA-seq quantification. Nat. Biotechnol. 34, 525–527 (2016).

74. M. I. Love, W. Huber, S. Anders, Moderated estimation of fold change and dispersion for RNA-seq data with DESeq2. Genome Biol. 15, 550 (2014).

75. A. Santos-Lopez, et al., Evolved resistance to a novel cationic peptide antibiotic requires high mutation supply. Evol. Med. Public Health 10, 266–276 (2022).

76. C. B. Turner, C. W. Marshall, V. S. Cooper, Parallel genetic adaptation across environments differing in mode of growth or resource availability. Evol. Lett. 2, 355–367 (2018).

77. D. E. Deatherage, J. E. Barrick, Identification of mutations in laboratory-evolved microbes from next-generation sequencing data using breseq. Methods Mol. Biol. 1151, 165–188 (2014).

78. A. Alonso-Del Valle, et al., Variability of plasmid fitness effects contributes to plasmid persistence in bacterial communities. Nat. Commun. 12, 2653 (2021).

79. M. J. Muraski, et al., Adaptation to Overflow Metabolism by Mutations That Impair tRNA Modification in Experimentally Evolved Bacteria. MBio 14, e0028723 (2023).

80. F. Lin, et al., Molecular Characterization of Reduced Susceptibility to Biocides in Clinical Isolates of Acinetobacter baumannii. Front. Microbiol. 8, 1836 (2017).

## Supplemental Methods Citations

1. C. D. M. Wijers, et al., Identification of Two Variants of Acinetobacter baumannii Strain ATCC 17978 with Distinct Genotypes and Phenotypes. Infect. Immun. 89, e0045421 (2021).

2. P. Baumann, M. Doudoroff, R. Y. Stanier, A study of the Moraxella group. II. Oxidative-negative species (genus Acinetobacter). J. Bacteriol. 95, 1520–1541 (1968).

3. A. C. Jacobs, et al., AB5075, a Highly Virulent Isolate of Acinetobacter baumannii, as a Model Strain for the Evaluation of Pathogenesis and Antimicrobial Treatments. MBio 5, e01076–14 (2014).

4. L. A. Gallagher, et al., Resources for Genetic and Genomic Analysis of Emerging Pathogen Acinetobacter baumannii. J. Bacteriol. 197, 2027–2035 (2015).

5. A. Santos-Lopez, C. W. Marshall, M. R. Scribner, D. J. Snyder, V. S. Cooper, Evolutionary pathways to antibiotic resistance are dependent upon environmental structure and bacterial lifestyle. eLife 8 (2019).

6. O. Schwengers, et al., Bakta: rapid and standardized annotation of bacterial genomes via alignment-free sequence identification. Microb. Genom. 7 (2021).

7. G. Tonkin-Hill, et al., Producing polished prokaryotic pangenomes with the Panaroo pipeline. Genome Biol. 21, 180 (2020).

8. L. Pritchard, R. H. Glover, S. Humphris, J. G. Elphinstone, I. K. Toth, Genomics and taxonomy in diagnostics for food security: soft-rotting enterobacterial plant pathogens. Anal. Methods 8, 12–24 (2016).

9. K. A. Jolley, M. C. J. Maiden, BIGSdb: Scalable analysis of bacterial genome variation at the population level. BMC Bioinformatics 11, 595 (2010).

10. M. Feldgarden, et al., AMRFinderPlus and the Reference Gene Catalog facilitate examination of the genomic links among antimicrobial resistance, stress response, and virulence. Sci. Rep. 11, 12728 (2021).

11. R. L. Tatusov, M. Y. Galperin, D. A. Natale, E. V. Koonin, The COG database: a tool for genome-scale analysis of protein functions and evolution. Nucleic Acids Res. 28, 33–36 (2000).

12. C. CLSI, M100: Performance Standards for Antimicrobial Susceptibility Testing, 32nd Ed. (Clinical and Laboratory Standards Institute, 2022).

13. A. C. Stephens, L. R. Thurlow, A. R. Richardson, Mechanisms Behind the Indirect Impact of Metabolic Regulators on Virulence Factor Production in Staphylococcus aureus. Microbiol. Spectr. 10, e0206322 (2022).

14. N. L. Bray, H. Pimentel, P. Melsted, L. Pachter, Near-optimal probabilistic RNA-seq quantification. Nat. Biotechnol. 34, 525–527 (2016).

15. M. I. Love, W. Huber, S. Anders, Moderated estimation of fold change and dispersion for RNA-seq data with DESeq2. Genome Biol. 15, 550 (2014).

16. T. Bliek, J. Chouaref, F. van der Kloet, J. Szkodon, M. Galland, Introductionto RNA-seq lesson (2020) (July 1, 2024).

17. A. Zhu, J. G. Ibrahim, M. I. Love, Heavy-tailed prior distributions for sequence count data: removing the noise and preserving large differences. Bioinformatics 35, 2084–2092 (2019).

18. M. R. Scribner, A. Santos-Lopez, C. W. Marshall, C. Deitrick, V. S. Cooper, Parallel Evolution of Tobramycin Resistance across Species and Environments. MBio 11 (2020).

19. A. Santos-Lopez, et al., Evolved resistance to a novel cationic peptide antibiotic requires high mutation supply. Evol. Med. Public Health 10, 266–276 (2022).

20. M. Baym, et al., Inexpensive multiplexed library preparation for megabase-sized genomes. PLoS ONE 10, e0128036 (2015).

21. C. B. Turner, C. W. Marshall, V. S. Cooper, Parallel genetic adaptation across environments differing in mode of growth or resource availability. Evol. Lett. 2, 355–367 (2018).

22. D. E. Deatherage, J. E. Barrick, Identification of mutations in laboratory-evolved microbes from next-generation sequencing data using breseq. Methods Mol. Biol. 1151, 165–188 (2014).

23. A. Alonso-Del Valle, et al., Variability of plasmid fitness effects contributes to plasmid persistence in bacterial communities. Nat. Commun. 12, 2653 (2021).

24. M. J. Muraski, et al., Adaptation to Overflow Metabolism by Mutations That Impair tRNA Modification in Experimentally Evolved Bacteria. MBio 14, e0028723 (2023).

25. S. Coyne, P. Courvalin, B. Périchon, Efflux-mediated antibiotic resistance in Acinetobacter spp. Antimicrob. Agents Chemother. 55, 947–953 (2011).

26. F. Lin, et al., Molecular Characterization of Reduced Susceptibility to Biocides in Clinical Isolates of Acinetobacter baumannii. Front. Microbiol. 8, 1836 (2017).

27. Z. R. Lonergan, et al., An Acinetobacter baumannii, Zinc-Regulated Peptidase Maintains Cell Wall Integrity during Immune-Mediated Nutrient Sequestration. Cell Rep. 26, 2009–2018.e6 (2019).

